# Prior induction of cellular antiviral pathways limits frog virus 3 replication in two permissive *Xenopus laevis* skin epithelial-like cell lines

**DOI:** 10.1101/2021.06.04.446995

**Authors:** Maxwell P. Bui-Marinos, Lauren A. Todd, Marie-Claire D. Wasson, Brandon E. E. Morningstar, Barbara A. Katzenback

## Abstract

Frog virus 3 (FV3) causes mortality in a range of amphibian species. Despite the importance of the skin epithelium as a first line of defence against FV3, the interaction between amphibian skin epithelial cells and FV3 remains largely uncharacterized. Here, we used newly established *Xenopus laevis* skin epithelial-like cell lines, Xela DS2 and Xela VS2, to study the susceptibility and permissiveness of frog skin epithelial cells to FV3, and the innate immune antiviral and proinflammatory gene regulatory responses of these cells to FV3. Both cell lines are susceptible and permissive to FV3, yet do not exhibit appreciable transcript levels of scavenger receptors recently demonstrated to be used by FV3 for cellular entry. Xela DS2 and Xela VS2 upregulate antiviral and proinflammatory cytokine transcripts in response to poly(I:C) but not to FV3 or UV-inactivated FV3. Poly(I:C) pretreatment limited FV3 replication and FV3-induced cytopathic effects in both cell lines. Thus, Xela DS2 and Xela VS2 can support FV3 propagation, represent *in vitro* systems to investigate antiviral responses of frog skin epithelial cells, and are novel tools for screening compounds that initiate effective antiviral programs to limit FV3 replication.

## 1. Introduction

*Frog virus 3* (FV3) is a large (105.9 kbp) double-stranded DNA virus and type species of the *Ranavirus* genus, family *Iridoviridae*. In North America, ranaviruses are known to infect at least 55 different amphibian species and have been implicated in mortality events in over 30 of these species (Miller et al., 2011). Susceptibility to FV3 differs across amphibian species (Hoverman et al., 2011) and within a species depending on the species’ life stage, genotype (Gantress et al., 2003), and environment (Brand et al., 2016). Susceptible amphibians exhibit severe pathology such as skin lesions, swelling, and internal hemorrhage (Forzán et al., 2017; Miller et al., 2007) that can result in up to 100% mortality in tadpoles (Haislip et al., 2011; Hoverman et al., 2011) and post-metamorphs (Forzán et al., 2015), but appears to vary with initial viral infection dose (Bienentreu et al., 2020). Relatively resistant amphibians develop mild symptoms including lethargy, skin shedding and cutaneous erythema, but often recover in 2 – 3 weeks (Gantress et al., 2003) and instead may serve as asymptomatic reservoirs of FV3 (Robert et al., 2007). Transmission of FV3 to naïve hosts can occur by water borne transmission or consumption of infected carcass (Harp and Petranka, 2006; Robert et al., 2011) and necessitates that FV3 evade a host epithelial barrier.

The African clawed frog (*Xenopus laevis*) was established as a model to study immune responses to FV3 in the early 2000s (Gantress et al., 2003), and research using this model has led to a basic understanding of the anuran anti-ranaviral immune response [reviewed in (Robert et al., 2017)]. FV3 infection of adult *X. laevis* results in an increase in type I interferon (IFN) and proinflammatory cytokine transcript levels in internal organs (De Jesús Andino et al., 2012; Grayfer et al., 2014, 2015a), along with the recruitment of macrophages to the site of intraperitoneal infection (De Jesús Andino et al., 2012). Type I IFN is critical for effective antiviral defences against FV3, as pretreatment of the A6 *X. laevis* kidney epithelial cell line or intraperitoneal injection of tadpoles with recombinant type I IFN prior to FV3 infection limited viral replication (Grayfer et al., 2014). While these earlier studies demonstrated the importance of amphibian type I IFN responses in antiviral defences, the use of intraperitoneal injection to establish consistent FV3 infections in frog hosts largely precluded assessment of the antiviral roles played by the skin barrier. However, as amphibian skin is in continuous interface with the external environment, it represents an important innate immune barrier and first line of defence against aquatic pathogens such as FV3 [reviewed in (Varga et al., 2019)]. More recent studies have performed *in vivo* FV3 infections through water bath exposure, which more closely approximates natural routes of FV3 transmission. These investigations have uncovered key roles for type I and type III IFN responses in the skin of FV3-challenged *X. laevis* adults and tadpoles, respectively (Wendel et al., 2017; Wendel et al., 2018), including the short-term protective effects of subcutaneous administration of type I IFN or type III IFN, albeit to a lesser extent, in highly susceptible *X. laevis* tadpoles against FV3 infection (Wendel et al., 2017).

While the above studies provide evidence that frog skin tissue contributes to antiviral defence against FV3, the precise roles of each cell type within the skin have yet to be determined. Skin tissue is complex and is comprised of many cell types, including epithelial and fibroblast cells. Upon infection, additional cell types (i.e. macrophages and granulocytes) are also recruited to the site of infection (Grayfer et al., 2015b). Macrophages appear to mediate antiviral responses in relatively FV3-resistant hosts (Grayfer and Robert, 2014, 2015), but the functions of other cell types in the anti-FV3 response require further investigation. The study of whole skin tissue gives us meaningful information regarding how cells function synergistically to respond to FV3, however, the complex cellular composition of skin tissue hinders our ability to elucidate the individual contribution of each cell type to antiviral defences. Skin epithelial cells are in constant direct contact with potential sources of FV3, yet the individual contribution of skin epithelial cells to anti-FV3 defences remains largely uncharacterized.

To better investigate the initial interaction between frog epithelial cells and FV3, we previously developed and characterized two skin epithelial-like cell lines from the dorsal and ventral skin tissues of *X. laevis*, named Xela DS2 and Xela VS2, respectively (Bui-Marinos et al., 2020). We have previously demonstrated that these cells can respond to poly(I:C), a synthetic analogue of viral double-stranded RNA and potent inducer of type I IFN responses, through the upregulation of antiviral and proinflammatory cytokine transcripts. In this study, we sought to (1) evaluate whether Xela DS2 and Xela VS2 are susceptible and permissive to FV3, (2) determine whether these cell lines initiate gene regulatory antiviral and proinflammatory responses to FV3, and (3) uncover whether prior establishment of antiviral programs in these cell lines would confer protection against FV3-associated cellular cytopathicity and limit viral replication.

## 2. Methods

### 2.1. Xela DS2 and Xela VS2 cell lines and media

The generation, characterization and maintenance of the *X. laevis* dorsal (Xela DS2) and ventral (Xela VS2) skin epithelial-like cell lines have been previously described by our research group (Bui-Marinos et al., 2020). Briefly, Xela cell lines were sub-cultured every 3 – 4 days at a 1:4 split and cultured at 26 °C in plug-seal tissue culture treated flasks (BioLite; Thermo Fisher Scientific) containing amphibian-adjusted Leibovitz’s L-15 (AL-15; Wisent Inc.) medium supplemented with 15% fetal bovine serum (FBS; VWR), herein referred to as Xela complete media. Adherent cells were detached using 0.175% trypsin, 1.55 mM EDTA solution that was prepared by diluting seven parts of 0.25% trypsin, 2.21 mM EDTA solution (Wisent Inc.) with three parts of sterile ultrapure water. Amphibian phosphate-buffered saline (APBS) was similarly prepared by diluting seven parts of sterile PBS with three parts of sterile ultrapure water. For all experiments, seeding densities were adjusted to account for the plating efficiencies (79% for Xela DS2 and 83% for Xela VS2). Since viral binding and infection efficiency is hindered at high FBS concentrations (Petricevich et al., 2001), experiments involving FV3 infection of Xela cell lines were performed using AL-15 supplemented with 2% FBS, herein referred to as Xela low serum media.

### 2.2. *Epithelioma Papulosum Cyprini* (EPC) cells

EPC cells were kindly provided by Dr. Niels C. Bols (University of Waterloo, Ontario, Canada) and were maintained at 26 °C in plug-seal tissue culture treated flasks containing Leibovitz L-15 supplemented with 10% FBS, herein referred to as EPC complete media. EPC cells were sub-cultured (1:4) every 5-7 days by washing with PBS and treatment with 0.25% trypsin, 2.21 mM EDTA (Wisent Inc.) to detach adhered cells. Cells were collected by centrifugation at 300 × *g* for 10 min and resuspended in fresh EPC complete media. EPC cells demonstrated 100% plating efficiency when enumerated the day after seeding. Although it is known that current EPC cell lineages are contaminated with fathead minnow cells, they are still deemed worth retaining as a cell line for the study of aquatic viruses (Winton et al., 2010). EPC cells were selected for our experiments since they have been widely used to propagate and study FV3 [e.g. (Ariel et al., 2009; Pham et al., 2015). For methods involving FV3 and EPC cells, Leibovitz’s L-15 supplemented with 2% FBS was used and will be referred to as EPC low serum media.

### 2.3. Propagation of FV3 in EPC cells

FV3 (Granoff strain, ATCC VR-567) was a kind gift from Dr. Niels C. Bols (University of Waterloo, Ontario, Canada). FV3 was propagated on EPC monolayers by adding 1 mL of stock FV3 to 9 mL of EPC low serum media, and the entire volume was incubated on a monolayer of EPC cells for 7 days at 26 °C. Seven days post-infection (dpi), virus-containing media from the flask of infected EPC cells was collected, underwent three freeze-thaw cycles at −80 °C, and was centrifuged at 1,000 × *g* for 10 min, filtered through a 0.22 μm PES filter (FroggaBio), and aliquoted into sterile 1.5 mL microfuge tubes for storage at −80 °C.

### 2.4. Determination of FV3 viral titres

The tissue culture infectious dose wherein 50% of cells are infected (TCID_50_/mL) values were determined for viral stocks and experimental samples using the Kärber method (Kärber, 1931), further modified by Pham and colleagues (Pham et al., 2011). Briefly, EPC cells were seeded in a 96-well tissue culture treated microwell plate (Thermo Fisher Scientific) at a final cell density of 100,000 cells/well in 0.1 mL EPC complete media and allowed to adhere overnight at 26 °C. The next day, the media was removed and 0.2 mL of a ten-fold dilution series of the sample to be tested (prepared in EPC low serum media) was applied to EPC monolayers. FV3-infected EPC cells were incubated for 10 d (determination of TCID_50_/mL values for viral stocks) or 7 d (determination of TCID_50_/mL values for experimental samples) at 26 °C prior to scoring for cytopathic effects (CPE). TCID_50_/mL values were multiplied by a factor of 0.7 to determine the approximate FV3 plaque forming unit (PFU)/mL values for multiplicity of infection (MOI) calculations (Knudson and Tinsley, 1974).

### 2.5. Resazurin and CFDA-AM assays

Cellular metabolic activity and membrane integrity were evaluated using a combined resazurin/5-carboxyfluorescein diacetate acetoxymethyl ester (CFDA-AM) assay (Dayeh et al., 2004). To assess the viability of both adherent and suspension cells present in a well, stock concentrations of resazurin (44 mM in APBS; Acros Organics) and CFDA-AM (800 μM in DMSO; Invitrogen) were diluted in APBS to working concentrations of 4,400 μM and 40 μM, respectively, before direct addition to the wells in a 1:10 ratio (final concentration of 440 μM resazurin and 4 μM CFDA-AM). Direct addition of resazurin and CFDA-AM to the culture media in each well permits viability assessment of both adherent and suspension cell populations. Plates were protected from light and allowed to incubate for 1 h at 26 °C. Fluorescence intensity was then measured for resazurin (535 nm/590 nm) and CFDA-AM (484 nm/530 nm) using a Cytation 5 multi-mode imaging plate reader (BioTek). In all cases, wells containing resazurin/CFDA-AM solution but without cells were included as a background control and their fluorescent values were subtracted from the values obtained from experimental samples.

### 2.6. Assessment of Xela DS2 and Xela VS2 susceptibility and permissibility to FV3

To assess the susceptibility of Xela DS2 (passages 63 – 66) and Xela VS2 (passages 68 – 72) to FV3, cells were seeded in a 48-well tissue culture treated plate (BioLite) at a final cell density of 50,000 cells/well in 0.3 mL Xela complete media and allowed to adhere overnight at 26 °C. The next day, media was removed, and cells were treated in triplicate with 0.25 mL of Xela low serum media alone (mock-infected control) or containing FV3 at MOIs of 0.0002, 0.002, 0.02, 0.2, 2 or 20. After 2 h of virus absorption at 26 °C, media was removed, and wells were washed three times with 300 μL of APBS prior to the addition of 500 μL of Xela low serum media. Plates were sealed with parafilm and incubated at 26 °C for the duration of the experiment. Total (adherent and suspension) cell viability was measured using the resazurin/CFDA-AM assay (Section 2.5) after 0, 1, 3, 5, 7, 10, and 14 dpi. Four independent trials were conducted (*n* = 4).

To assess the permissibility of Xela DS2 (passages 129 – 139) and Xela VS2 (passages 134 – 139) to FV3 replication, 250,000 cells/well were seeded in a 12-well plate (Thermo Fisher Scientific) in 0.75 mL Xela complete media and allowed to adhere overnight at 26 °C. The next day, media was removed from the wells and cells were treated with 0.5 mL of Xela low serum media alone (mock-infected control) or containing FV3 at a MOI of 0.002, 0.02, 0.2, 2 or 20. After 2 h of virus absorption at 26 °C, media was removed and wells were washed three times with 0.5 mL of APBS, prior to the addition of 2.5 mL of fresh Xela low serum media to all wells. Plates were sealed with parafilm and incubated at 26 °C. Phase contrast digital images and cell culture media were collected on 0 (30 min post-wash), 1, 3, 5, and 7 dpi for all treatments. Phase contrast digital images were captured using a Leica DMi1 microscope fitted with a MC170 color camera and LAS X 4.8 software. Collected media was centrifuged at 500 × *g* for 5 min, and supernatants were transferred to sterile 1.5 mL microfuge tubes prior to storage at −80 °C. TCID_50_/mL values were determined as described in Section 2.4. This experiment was conducted four independent times (*n* = 4).

### 2.7. Detection of scavenger receptor transcripts

Total RNA was isolated from 3 **×** 10^6^ Xela DS2 (passages 28, 29 and 31) and Xela VS2 (passages 42, 46 and 48) using TRI reagent (Invitrogen) according to the manufacturer’s instructions with the following modifications: ground tissue samples were further homogenized using a 1 mL syringe fitted with a 25-gauge needle and RNA pellets were washed with 1 mL of 75% ethanol (twice for cell lines and five times for tissues). RNA was quantified using a NanoDrop 2000 spectrophotometer, and RNA quality was examined by electrophoresing 1 μg of RNA on a 1% agarose gel containing 1% bleach (Aranda et al., 2012) and 1 × RedSafe nucleic acid staining solution (FroggaBio) in 1 × TAE buffer at 100 V for 35 min. RNA was stored at −80 °C until use.

RNA (1 μg) was treated with 0.5 U of DNase I (Thermo Fisher Scientific), DNase I was heat-inactivated, and RNA was reverse-transcribed into cDNA using the 5 × All-In-One RT Master Mix (Bio Basic) according to the manufacturer’s specifications (25 °C for 10 min, 42 °C for 50 min and 85 °C for 5 min). Genomic DNA contamination was excluded by employing a no reverse-transcriptase (-RT) control. Synthesized cDNA was stored at −20 °C until use. Subsequent PCR reactions were set up with the GeneDireX kit [1 × reaction buffer with 2 mM MgCl_2_, 200 μM dNTPs, 200 nM (each) forward and reverse primers (Supplementary Table 1), 0.625 U *Taq* polymerase and 4 μL of 1:8 diluted cDNA]. Thermocycling conditions were as follows: 95 °C for 5 min, followed by 26 (*actb* or 35 (scavenger receptors) cycles of 95 °C for 30 sec, 55 °C (*actb*) or 52-54 °C (scavenger receptors) for 30 s and 72 °C for 30 s, and a final extension at 72 °C for 10 min. One-third of the PCR volume was visualized on a 1% agarose gel run in 1 × TAE and imaged using a ChemiDoc imager (Bio-Rad). We validated each amplicon by direct sequencing at The Centre for Applied Genomics (The Hospital for Sick Children) and confirmed amplicon identity by BLASTn analysis (Altschul et al., 1990).

### 2.8. UV inactivation of FV3

FV3 aliquots were thawed and gently vortexed prior to exposure to 150 mJ UV energy using a UV Crosslinker (BioRad), a dose which has been previously used to effectively inactivate FV3 (Chinchar et al., 2003). FV3 inactivation was confirmed by measuring levels of the viral *mcp* transcript (Supplementary Methods Section 5) and assessment of viral titres (Section 2.4). UV-irradiated FV3 aliquots were stored at −80 °C until use.

### 2.9. Challenge of Xela DS2 and Xela VS2 with FV3

Xela DS2 (passages 120 – 130) and Xela VS2 (passages 125 – 135) (*n* = 4 independent trials) were seeded in a 6-well plate (Eppendorf) at a cell density of 625,000 cells/well in 1 mL of Xela complete media and allowed to adhere overnight at 26 °C. The next day, the media was removed, and cells were treated with 0.5 mL of Xela low serum media alone (mock-infected control), UV-inactivated FV3 (MOI 2), or FV3 (MOI 2). After 2 h incubation at 26 °C, the media was removed, wells were washed three times with 0.5 mL of APBS, and 2 mL of fresh Xela low serum media was added to each well. To one of the monolayers previously incubated for 2 h with Xela low serum media alone, Xela low serum media containing 1 μg/mL poly(I:C) was added (positive control). Phase-contrast images were taken of all treatments at 0, 6, 24, 48, and 72 h post-treatment (commenced following the addition of fresh Xela low serum media) using a Leica DMi1 microscope fitted with a MC170 color camera and LAS X 4.8 software to assess cell morphology. At each time point, media was collected and centrifuged at 500 × *g* for 5 min. Cleared media was transferred to sterile 1.5 mL microfuge tubes and stored at −80 °C for determination of viral titres (see Section 2.4). Total RNA was isolated from combined adherent and suspension cells using the EZ-10 Spin Column Total RNA Minipreps Super Kit (Bio Basic) with the following modification: cells pelleted from culture media were suspended in 100 μL lysis solution then mixed 1:1 with 70% ethanol and stored on ice. Meanwhile, remaining adherent cells were washed once with 1 mL APBS, followed by the addition of 300 μL lysis solution directly to wells for 1 min on ice, then mixed 1:1 with 70% ethanol. Lysates from the suspension and adherent cells were combined and mixed by inversion prior to addition to spin columns. Total RNA isolation, cDNA synthesis, and reverse transcriptase-quantitative polymerase chain reaction (RT-qPCR) were performed as described in Section 2.10 and Section 2.11.

### 2.10. Total RNA isolation and cDNA synthesis for antiviral gene expression assays

Total RNA was isolated from Xela DS2, Xela VS2 and EPC cells using the EZ-10 Spin Column Total RNA Minipreps Super Kit (Bio Basic) according to the manufacturer’s specifications with modifications to include an on-column DNase I digestion as described previously by our group (Bui-Marinos et al., 2020). RNA quantity and purity were determined as described in Section 2.7. RNA (500 ng) was reverse-transcribed into cDNA using the SensiFAST cDNA Synthesis Kit (BioLine) according to the manufacturer’s specifications for synthesis of cDNA for use in RT-qPCR reactions. Synthesized cDNA was stored at −20 °C until use.

### 2.11. RT-qPCR

Primer sequences, accession numbers, R^2^ and primer efficiency values for *X. laevis tnf, il1b, cxcl8a, ikb, ifn, mx2, pkr, actb, cyp, ef1a, gapdh*, and *hgprt* targets used in this study have been previously reported (Bui-Marinos et al., 2020). The type I IFN (*ifn*) primers targets a conserved region of *ifn3, ifn4, ifn6 and ifn7* from the expanded family of *X. laevis* type I IFNs (Bui-Marinos et al., 2020). Gene stability measures (M-values) were determined for all endogenous control candidates (*actb, cyp, ef1a, gapdh*, and *hgprt;* Supplementary Table 2) to identify a suitable endogenous control, as described in (Bui-Marinos et al., 2020), using the Applied Biosystems QuantStudio Analysis software, wherein a lower M-value infers stronger gene stability across time and treatment. With M-values of 0.428 and 0.424 for Xela DS2 and Xela VS2 samples, respectively, *actb* was selected as the endogenous reference gene for analysis of relative transcript abundance.

For relative transcript abundance analysis, RT-qPCR reactions were prepared in duplicate and consisted of 2.5 μL of 500 nM sense and antisense primers, 5 μL PowerUp SYBR green mix (Thermo Fisher Scientific), and 2.5 μL of diluted (1:20) cDNA template. Thermocycling conditions were as follows: initial denaturation at 50 °C for 2 min, followed by 95 °C for 2 min and 40 amplification cycles of denaturation at 95 °C for 1 s and extension at 60 °C for 30 s. A melt curve step followed all runs to ensure only a single dissociation peak was present, with initial denaturation at 95 °C for 1 s, then dissociation analysis at 60 °C for 20 s followed by 0.1 °C increments between 60 °C and 95 °C at 0.1 °C/s. Reactions were prepared in MicroAmp fast optical 96-well reaction plates (Life Technologies), sealed with MicroAmp clear optical film (Life Technologies) and run on a QuantStudio5 Real-Time PCR System (Thermo Fisher Scientific).

### 2.12. Effect of poly(I:C) treatment on Xela DS2 and Xela VS2 cell viability

Xela DS2 (passages 33 – 35) and Xela VS2 (passages 41 – 43) cells were seeded in a 48-well plate at a final cell density of 50,000 cells/well in 0.3 mL Xela complete media and allowed to adhere overnight at 26 °C. The next day, media was removed, and cells were treated with 500 μL of Xela low serum media containing 0, 10, 50, 100, 1,000, or 10,000 ng/mL poly(I:C) and incubated at 26 °C. Resazurin/CFDA-AM assays were performed as previously described (Section 2.5) after 1, 2, 3, and 5 days post-treatment to measure total cell (adherent and suspension) viability. Three independent trials were performed for each set of experiments (*n* = 3).

### 2.13. Antiviral assays

Xela DS2 (passages 135 – 140) and Xela VS2 (passages 138 – 143) cells were seeded in a 48-well plate (Thermo Fisher Scientific) at a final cell density of 50,000 cells/well in 0.3 mL Xela complete media and allowed to adhere overnight at 26 °C. The next day, media was removed, and cells were pre-treated with 160 μL of Xela low serum media containing 0, 10, 50, or 100 ng/mL poly(I:C) for 24 h at 26 °C, in sextuplicate. Following pretreatment, 90 μL of Xela low serum media alone was added to three of the six replicates for all treatments (mock-infected), and Xela low serum media containing FV3 (MOI 2) was added to the remaining triplicate pre-treated wells. After incubation at 26 °C for 2 h, media was removed from all wells which were then washed three times with 0.3 mL of APBS prior to the addition of 0.5 mL of fresh Xela low serum media and subsequent incubation at 26 °C. After 0, 3, and 5 dpi, cell culture media was collected, cleared by centrifugation at 500 × *g* for 5 min, and transferred to sterile 1.5 mL microfuge tubes prior to storage at −80 °C for future determination of TCID_50_/mL values (see Section 2.4). Afterwards, duplicate wells were treated with 200 μL of 440 μM resazurin solution dissolved in APBS to determine adherent cell metabolic activity, and a single well was treated with 200 μL Xela low serum media containing NucBlue Live reagent (Thermo Fisher Scientific). Following 1 h of incubation in the dark, plates were read on a BioTek Cytation 5 multimode plate reader and imager using Gen5 software under the following conditions: resazurin-treated wells were read with an excitation wavelength of 535 nm and emission wavelength 590 nm, while the NucBlue Live-treated wells were fluorescently imaged with an excitation wavelength of 377 nm and emission wavelength 477 nm to determine representative cell counts for each respective treatment. Digital phase contrast images were captured in addition to fluorescent imaging of NucBlue Live-treated cells. Four independent trials were conducted, using cells from different passages for each trial (*n* = 4).

### 2.14. Statistics

Prior to all statistical analyses, datasets were tested for normality using the Shapiro-Wilks test. Resazurin/CFDA-AM assay data for cellular viability after FV3 exposure (Section 2.6) and poly(I:C) cytotoxicity (Section 2.12) data were analyzed using a two-way ANOVA test followed by a Tukey’s post-hoc test, or a Kruskal-Wallis test followed by Dunn’s post-hoc test in cases where data was not normally distributed (Supplementary Figure 1B). RT-qPCR (Section 2.11) data were analyzed using a Kruskal-Wallis test followed by Dunn’s post-hoc test. Data regarding the effect of poly(I:C) pretreatment on FV3 replication based on cellular adherence and FV3 titres (Section 2.13) were analyzed using a two-way ANOVA test followed by a Tukey’s post-hoc test. All statistical analyses were performed using GraphPad Prism v8 software and groups were considered statistically significant when *p* < 0.05.

## 3. Results

### 3.1. Xela DS2 and Xela VS2 are susceptible and permissive to FV3

The susceptibility of Xela DS2 and Xela VS2 to FV3 (MOI 0.002 – 20) was assessed over 14 dpi using a combined resazurin/CFDA-AM assay to monitor changes in cell metabolic activity and cell membrane integrity of the entire cell population. We observed FV3 MOI to have a significant effect on the metabolic activity of Xela DS2 (Fig. 1A) and Xela VS2 (Fig. 1B) (two-way ANOVA, *p* < 0.0001), along with a significant interaction between MOI and time (two-way ANOVA, Xela DS2, *p* = 0.0024; Xela VS2, *p* = 0.0002). At a MOI of 20, a significant decrease in cell metabolic activity was observed for Xela DS2 (Fig. 1A) and Xela VS2 (Fig. 1B) as early as 3 dpi, and cell metabolic activity was virtually undetectable by 7 dpi. A significant decrease in metabolic activity of Xela DS2 (Fig. 1A) and Xela VS2 (Fig. 1B) was also observed as early as 5 dpi at a MOI of 2. Although not statistically significant, a decrease in metabolic activity of Xela DS2 and Xela VS2 was also observed at 7 dpi at a MOI of 0.2 (Fig 1A, B, respectively). A similar MOI-dependent decrease in cell membrane integrity was observed for Xela DS2 (Fig. 1C) and Xela VS2 (Fig. 1D) (two-way ANOVA, *p* < 0.0001), as well a significant interaction between MOI and time (two-way ANOVA, Xela DS2, *p* = 0.0011; Xela VS2, *p* = 0.0348). No statistically significant differences were observed in relation to Xela DS2 or Xela VS2 metabolic activity or membrane integrity between mock-infected cells and FV3-infected cells at a MOI of 0.02 or lower over 7 dpi (Fig. 1). A similar trend was observed at 14 dpi for both cell lines, although metabolic activity and membrane integrity were reduced and somewhat variable by 14 dpi for both cell lines across all treatment groups (Fig. 1). FV3-infected Xela DS2 and Xela VS2 monolayers (passages 129 – 139) exhibited dose- and time-dependent CPE (Fig. 2A, B) that preceded the loss of cell viability (Fig. 1). CPE were characterized by cell contraction and loss of adherence to the cell culture vessel, yielding floating cell clusters in the culture media and culminating in complete destruction of the monolayer. While Xela DS2 (Fig. 2A) and Xela VS2 (Fig. 2B) exhibited CPE when infected with FV3 at higher MOI (0.02 – 20), CPE was not observed in monolayers infected with FV3 at a MOI of 0.002. Similar dose- and time-dependent changes in cell metabolic activity (Supplementary Figure 1B) and morphology (Supplementary Figure 1A) were also observed in earlier passages (passage 20 – 55) of Xela DS2 and Xela VS2 infected with FV3.

**Figure 1.**
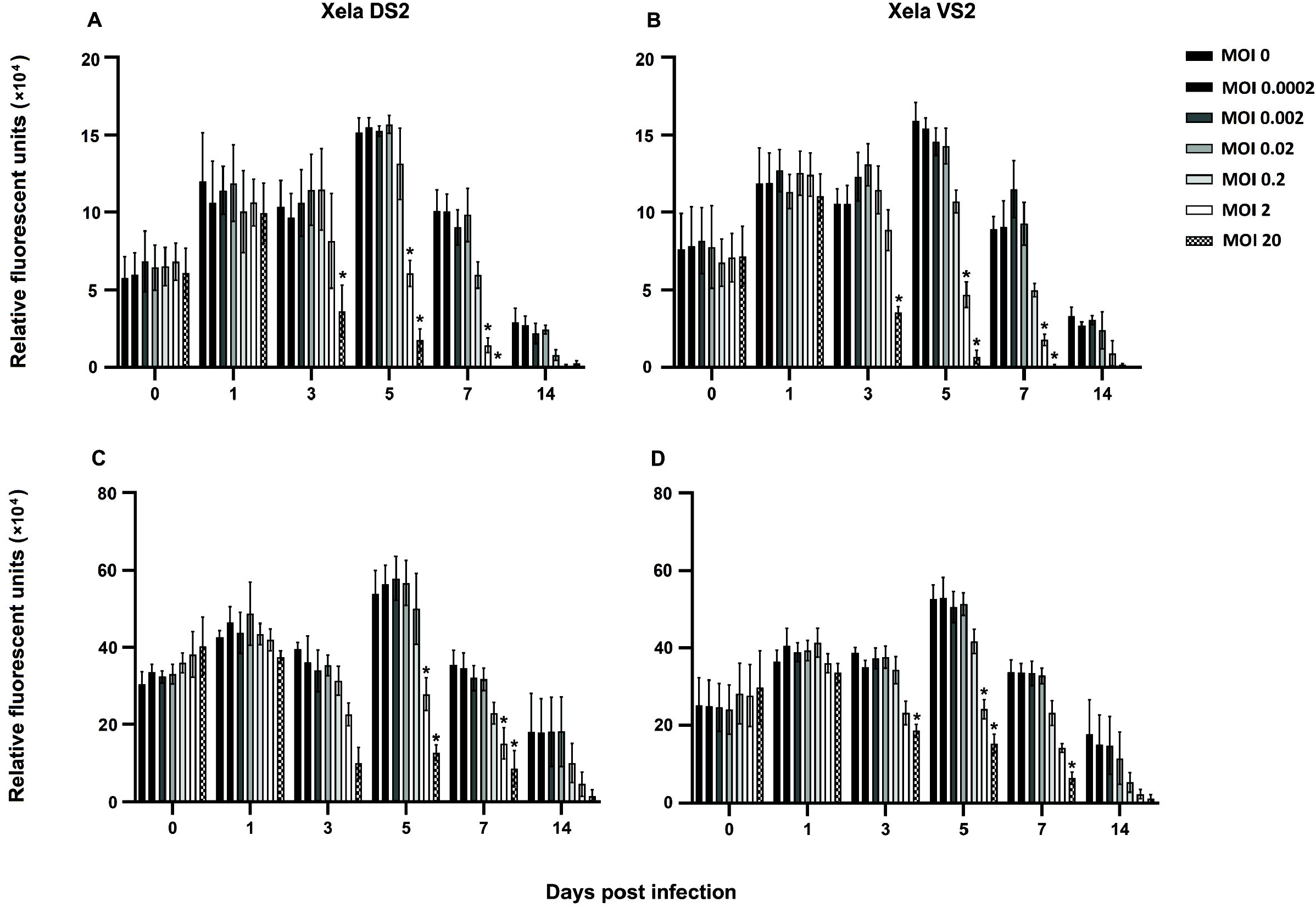
Xela DS2 and Xela VS2 are susceptible to FV3 and exhibit a loss of cell viability at high MOIs. Xela DS2 and Xela VS2 cells were infected with FV3 by absorption for 2 h at a MOI of 0 (mock-infected), 0.0002, 0.002, 0.02, 0.2, 2 or 20. Over 14 dpi, (A) Xela DS2 and (B) Xela VS2 cell metabolic activity (rezazurin assay) and (C) Xela DS2 and (D) Xela VS2 membrane integrity (CFDA-AM assay) were measured on a BioTek Cytation 5 multimode plate reader and imager. Data represent the mean relative fluorescent units ± standard error and were analyzed with a two-way ANOVA with Tukey’s post-hoc test (*p* < 0.05). *n* = 4 independent experiments. Asterisks (*) indicate statistical significance of the infected cells relative to mock-infected cells (MOI 0) within each respective day.

**Figure 2.**
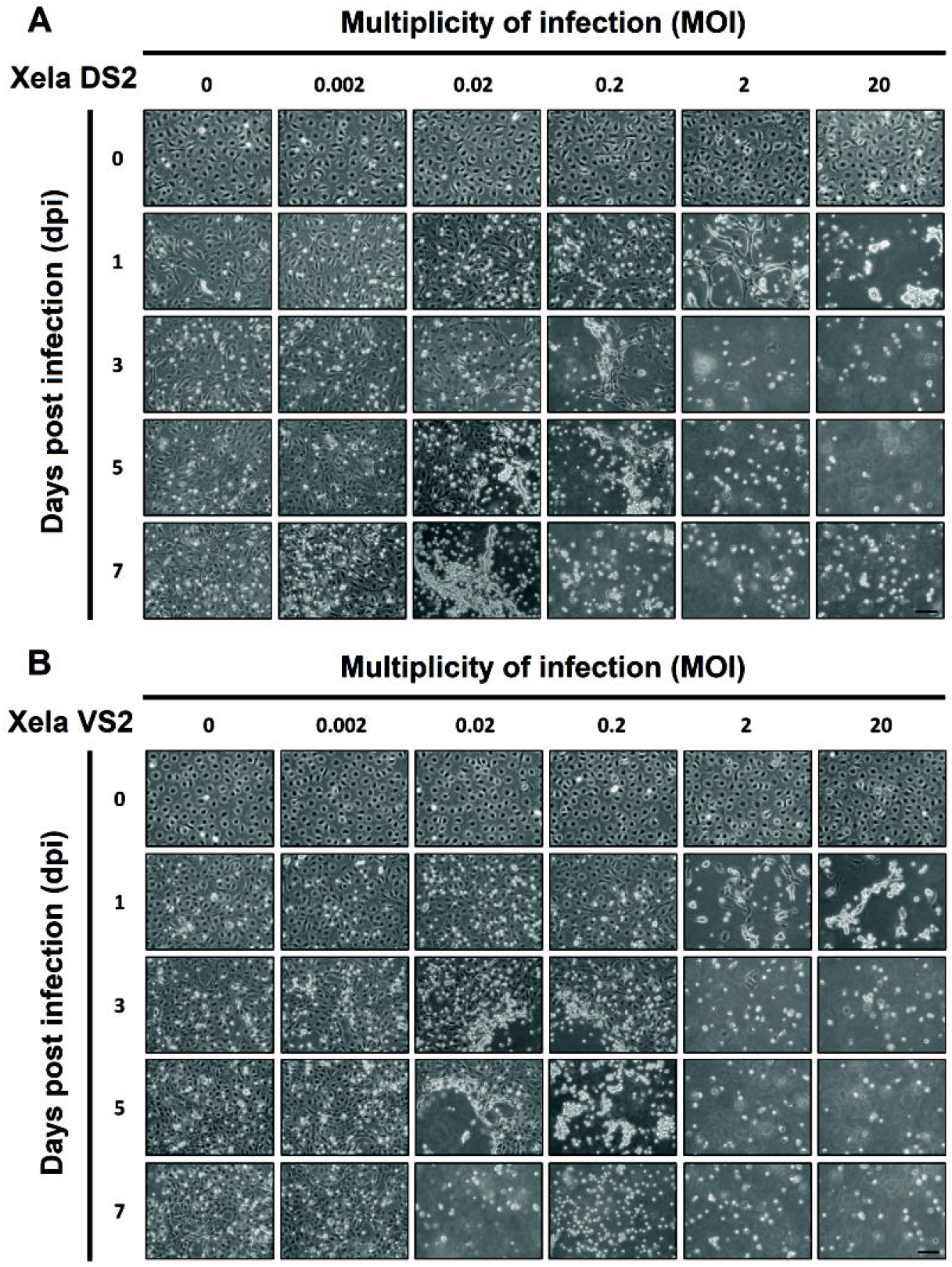
Cytopathic effects are observed in FV3-infected Xela DS2 and Xela VS2. Monolayers of (A) Xela DS2 and (B) Xela VS2 were infected with FV3 by absorption for 2 h at a MOI of 0 (mock-infected), 0.0002, 0.002, 0.02, 0.2, 2 or 20 and cell morphology was documented over 7 dpi. Phase contrast images were captured at 200 × magnification using a Leica DMi1 microscope (scale bar is 100 μM). The images shown are representative of four independent trials.

To assess whether Xela DS2 and Xela VS2 are permissive to FV3, monolayers were mock-infected or infected with FV3 at MOIs of 0.002 – 20, and viral titres were measured over 7 dpi by determining TCID_50_/mL values. Xela DS2 (Fig. 3A) and Xela VS2 (Fig. 3B) infected with FV3 at a MOI of 0.02 or greater demonstrated marked increases in viral titres in a dose- and time-dependent manner, with viral titre levels peaking at 5-7 dpi [maximal log_10_(TCID_50_/mL) value of 6-7]. While Xela DS2 and Xela VS2 supported FV3 replication when infected at higher MOIs (0.02 – 20), FV3 appeared unable to effectively replicate in Xela DS2 and Xela VS2 at a MOI of 0.002 over the period examined (Fig. 3A, B). No viral particles were detected in culture media collected from mock-infected (MOI 0) Xela DS2 or Xela VS2 at any time point (Fig. 3). Similar time- and dose-dependent FV3 replication was observed in earlier passages (passage 30 – 50) of Xela DS2 and Xela VS2 as determined through viral titres (Supplementary Figure 1C) and detection of viral transcripts (Supplementary Figure 2).

**Figure 3.**
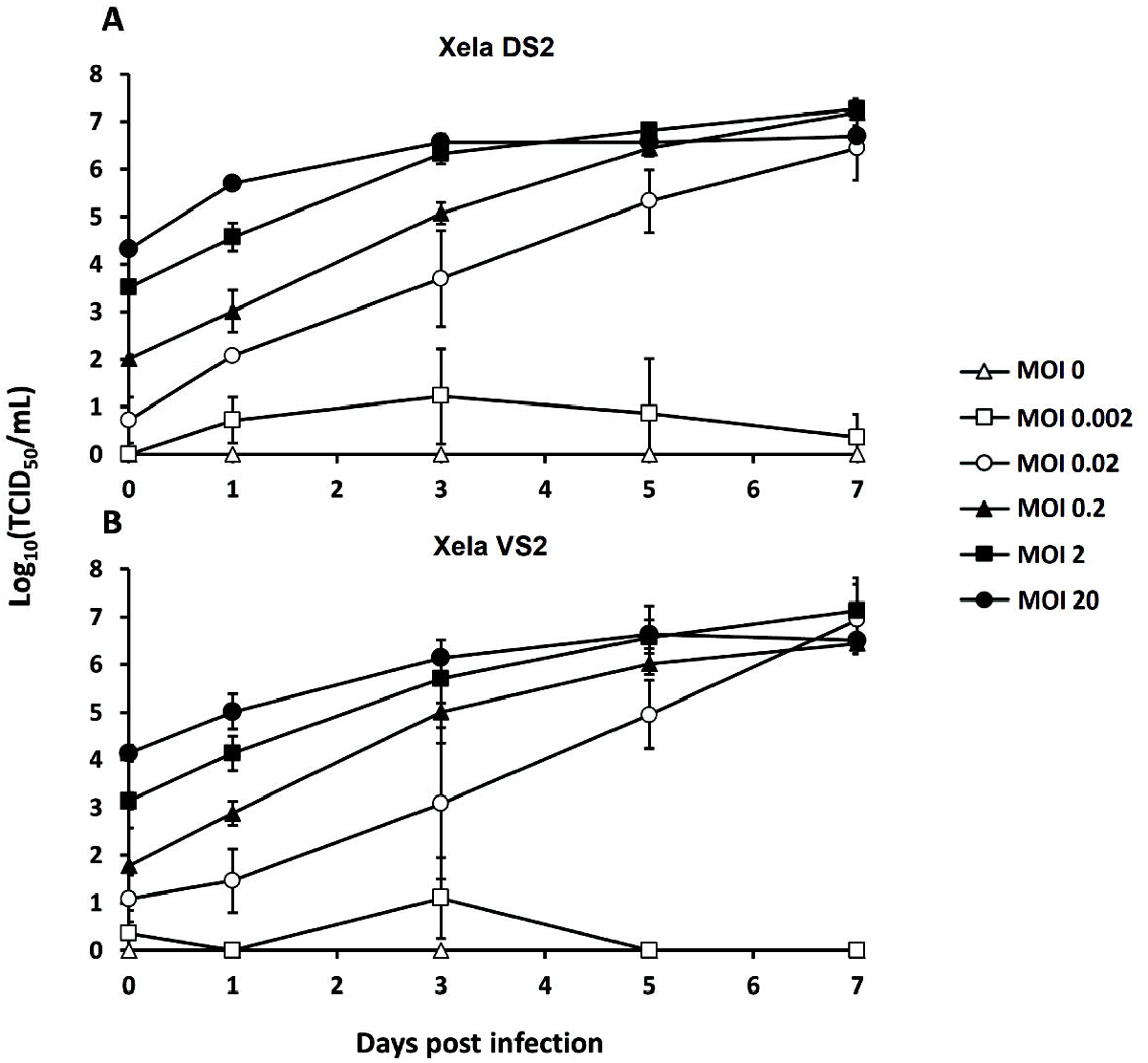
Xela DS2 and Xela VS2 are permissive to FV3. Monolayers of (A) Xela DS2 (passage 129 – 139) and (B) Xela VS2 (passage 134 – 139) were infected with FV3 by absorption for 2 h at a MOI of 0 (mock-infected), 0.0002, 0.002, 0.02, 0.2, 2 or 20. Monolayers were washed three times with APBS to remove non-absorbed virus, and low serum media was added to the monolayers. Virus-containing media was removed 30 min after addition (day 0) and at 1, 3, 5 and 7 dpi. Samples were serially diluted and applied to EPC monolayers and wells were scored for CPE after 7 dpi to determine TCID_50_/mL values. CPE were not observed in EPC monolayers treated with cell culture media collected from mock-infected (A) Xela DS2 or (B) Xela VS2 at any time point. Data were log10 transformed and are presented as the mean ± standard error of four independent experiments.

### 3.2. Xela DS2 and VS2 do not express appreciable levels of class A scavenger receptors thought to be used by FV3 during cellular entry

Given that Xela DS2 and Xela VS2 are susceptible to FV3 infection, we sought to examine the expression of class A scavenger receptors, a class of proteins that have been shown to be used by FV3 during viral entry (Vo et al., 2019a). We examined the expression of transcripts corresponding to class A scavenger receptors in Xela DS2 and Xela VS2. RT-PCR analyses revealed a general lack of class A scavenger receptor (*srai/ii, scara3, scara4, scara5, marco*) transcripts in Xela DS2 and Xela VS2, despite the detection of abundant levels in reference tissues (Fig. 4). *scara3*, *scara4* and *scara5* transcripts were detected in control spleen tissue as well as dorsal and ventral skin tissue, while *srai/ii* and *marco* transcripts were detected in control spleen tissue but were undetected in dorsal and ventral skin tissue. Very low levels of *scara3* and *scara5* transcripts were detected in two passages of Xela DS2 and Xela VS2 cells, respectively.

**Figure 4.**
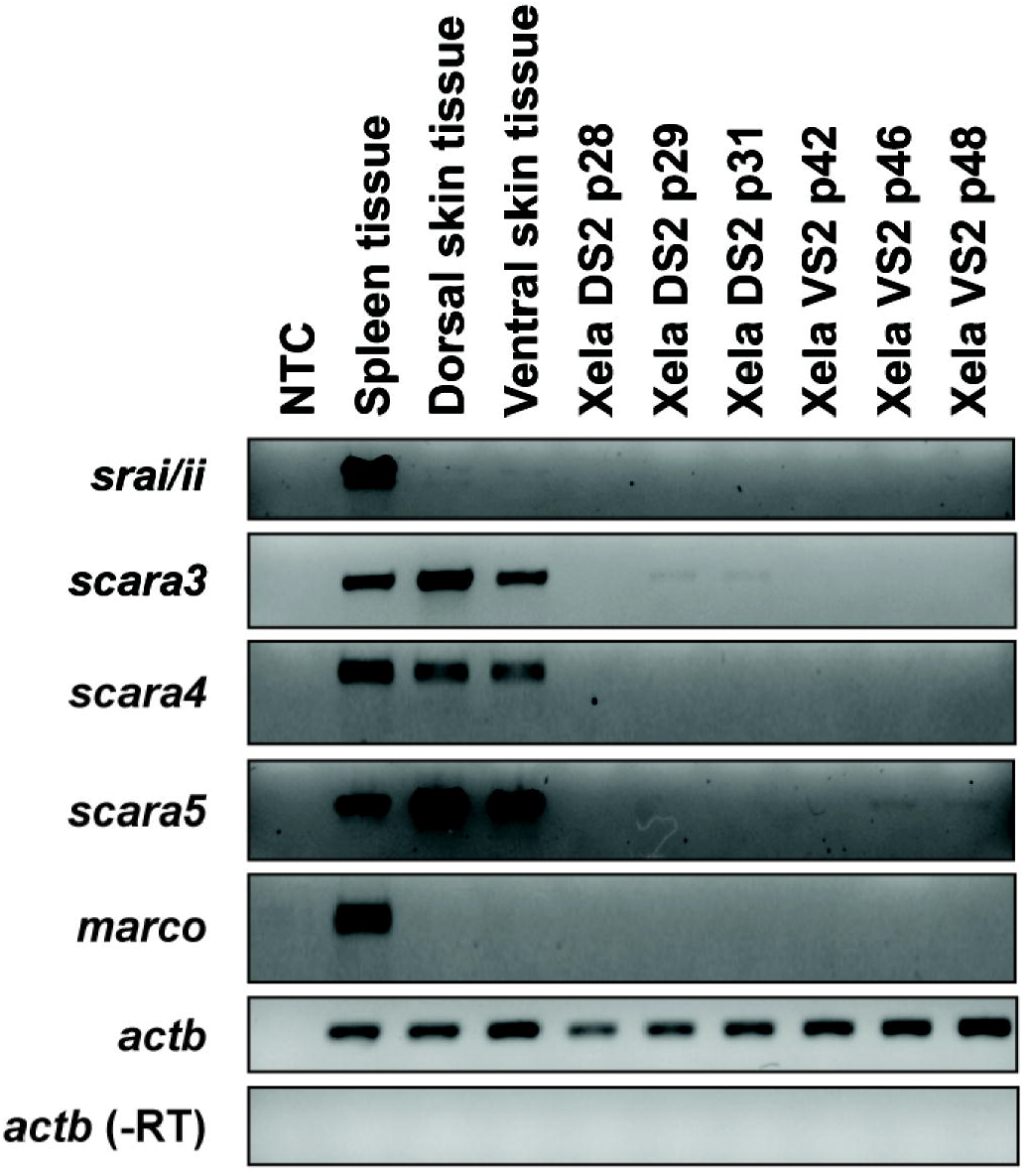
Transcripts corresponding to known *X. laevis* class A scavenger receptors are virtually undetectable in Xela DS2 and Xela VS2. Total RNA was isolated from Xela DS2 (passages 28, 29 and 31) and Xela VS2 (passages (42, 46 and 48), as well as control tissues (spleen, dorsal skin and ventral skin). RT-PCR targets included *X. laevis srai/ii, scara3, scara4, scara5* and *marco* transcripts. Amplification of *actb* served as an endogenous control, while amplification of *actb* in cDNA samples prepared without reverse-transcriptase (-RT) indicated the absence of contaminating genomic DNA.

### 3.3. Xela DS2 and Xela VS2 do not upregulate antiviral or proinflammatory transcripts when challenged with UV-inactivated FV3 or FV3

We previously demonstrated that Xela DS2 and Xela VS2 upregulated key antiviral and proinflammatory transcripts in response to poly(I:C), indicating that Xela DS2 and Xela VS2 are capable of activating intrinsic antiviral pathways following recognition of a synthetic analogue of viral double-stranded RNA (Bui-Marinos et al., 2020). To determine whether Xela DS2 and Xela VS2 mount antiviral and proinflammatory responses to FV3 at the transcriptional level, Xela DS2 and Xela VS2 were challenged with 1 μg/mL poly(I:C), UV-inactivated FV3 at a MOI of 2, or FV3 at a MOI of 2, and mRNA levels of antiviral (*ifn, mx2*, and *pkr*), pro-inflammatory (*il1b, tnf*, and *cxcl8*) and *ikb* genes were measured using RT-qPCR. Similar to our previous study (Bui-Marinos et al., 2020), poly(I:C) treatment of Xela DS2 and Xela VS2 resulted in significant increases in *pkr* (Fig. 5E, F), *il1b* (Fig. 6A, B), *tnf* (Fig. 6C, D), *cxcl8* (Fig. 6E, F) and *ikb* (Fig. 6G, H) transcript levels in comparison to the time-matched mock-treated cells (media alone) over the 72 h examined, whereas no statistically significant changes in *ifn* (Fig. 5A, B) or *mx2* (Fig. 5C, D) transcript levels were observed at any time point. In contrast, challenge of Xela DS2 or Xela VS2 with UV-inactivated FV3 or FV3 did not appear to induce the expression of the antiviral (Fig. 5) or proinflammatory (Fig. 6) gene targets examined over the 72-h period, despite observed changes in cell morphology that resulted in loss of cellular adherence to the tissue culture vessel (Supplementary Figure 3). Although CPE were observed in Xela DS2 and Xela VS2 challenged with UV-inactivated FV3 (MOI 2), transcripts for the FV3 *mcp* gene were undetectable (Supplementary Figure 4A) and TCID_50_/mL values did not increase over time (Supplementary Fig. 4B, C), confirming that UV-inactivated FV3 could not undergo viral replication. In contrast to these results, we detected *mcp* transcripts (Supplementary Figure 4A) and observed increasing TCID_50_/mL values in FV3-infected Xela DS2 and Xela VS2 (Supplementary Figure 4B, C).

**Figure 5.**
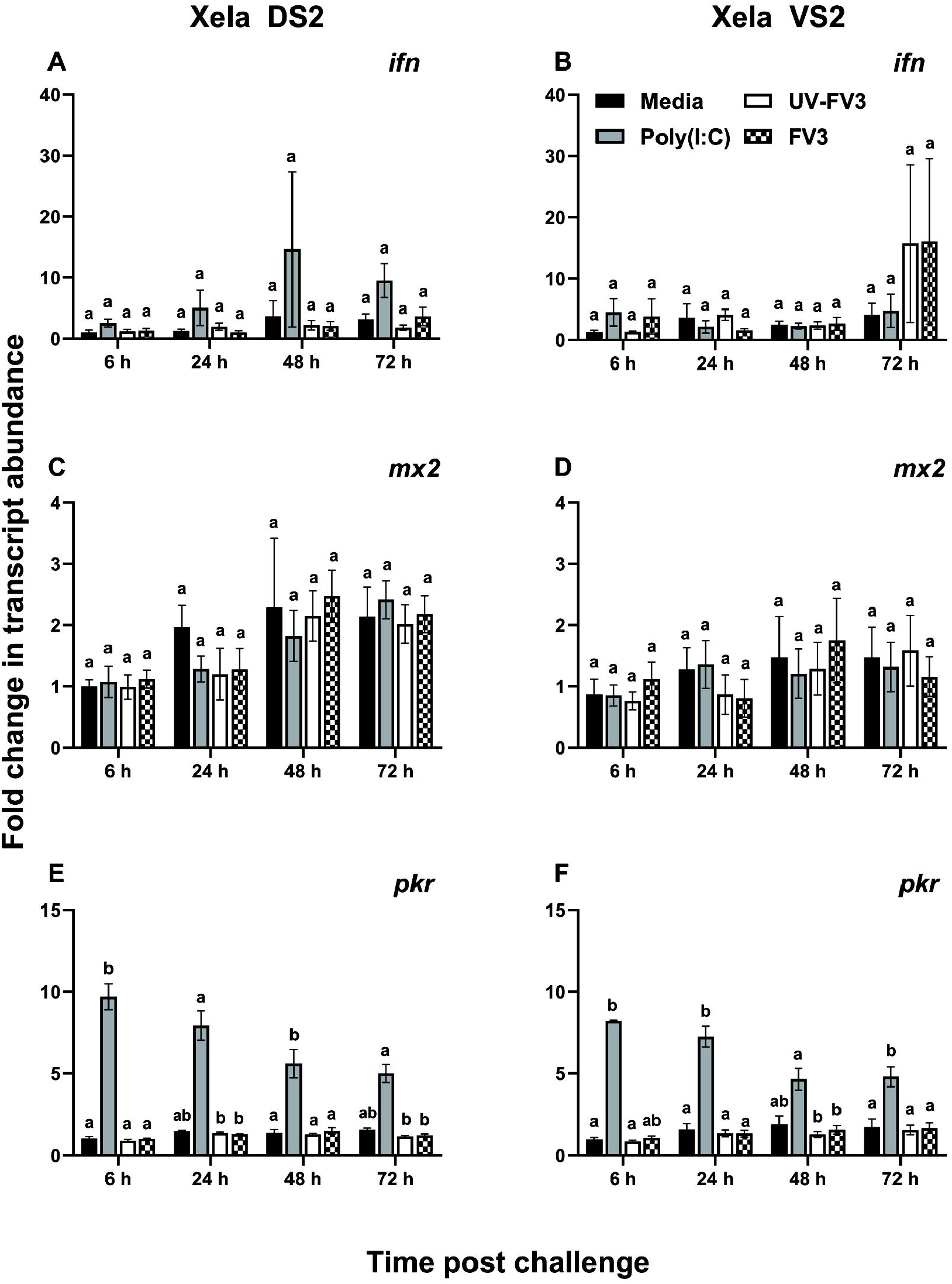
Xela DS2 and Xela VS2 do not upregulate antiviral transcripts following challenge with UV-inactivated FV3 or FV3. Xela DS2 or Xela VS2 were mock-infected (media), treated with 1 μg/mL poly I:C (pIC), challenged with UV-inactivated FV3 at a MOI of 2 or infected with FV3 at a MOI of 2 for 6 h, 24 h, 48 h or 72h. RT-qPCR was performed using cDNA generated from each treatment and time point to determine relative transcript levels of (A,B) *ifn*, (C,D) *mx2*, and (E,F) *pkr*. The type I IFN primer set (*ifn*) targets a highly conserved region of *X. laevis ifn3, ifn4, ifn6 and ifn7*. Data were analyzed using the ΔΔCt method and were expressed as a fold-change in transcript levels relative to that of the 6 h non-treated control sample. Data represent the mean ± standard error and were analyzed with a Kruskal-Wallis test and Dunn’s post-hoc test. *n* = 4 independent experiments. Within each time point, significant statistical differences (*p* < 0.05) are denoted using a lettering system, wherein groups with the same lettering are not statistically different.

**Figure 6.**
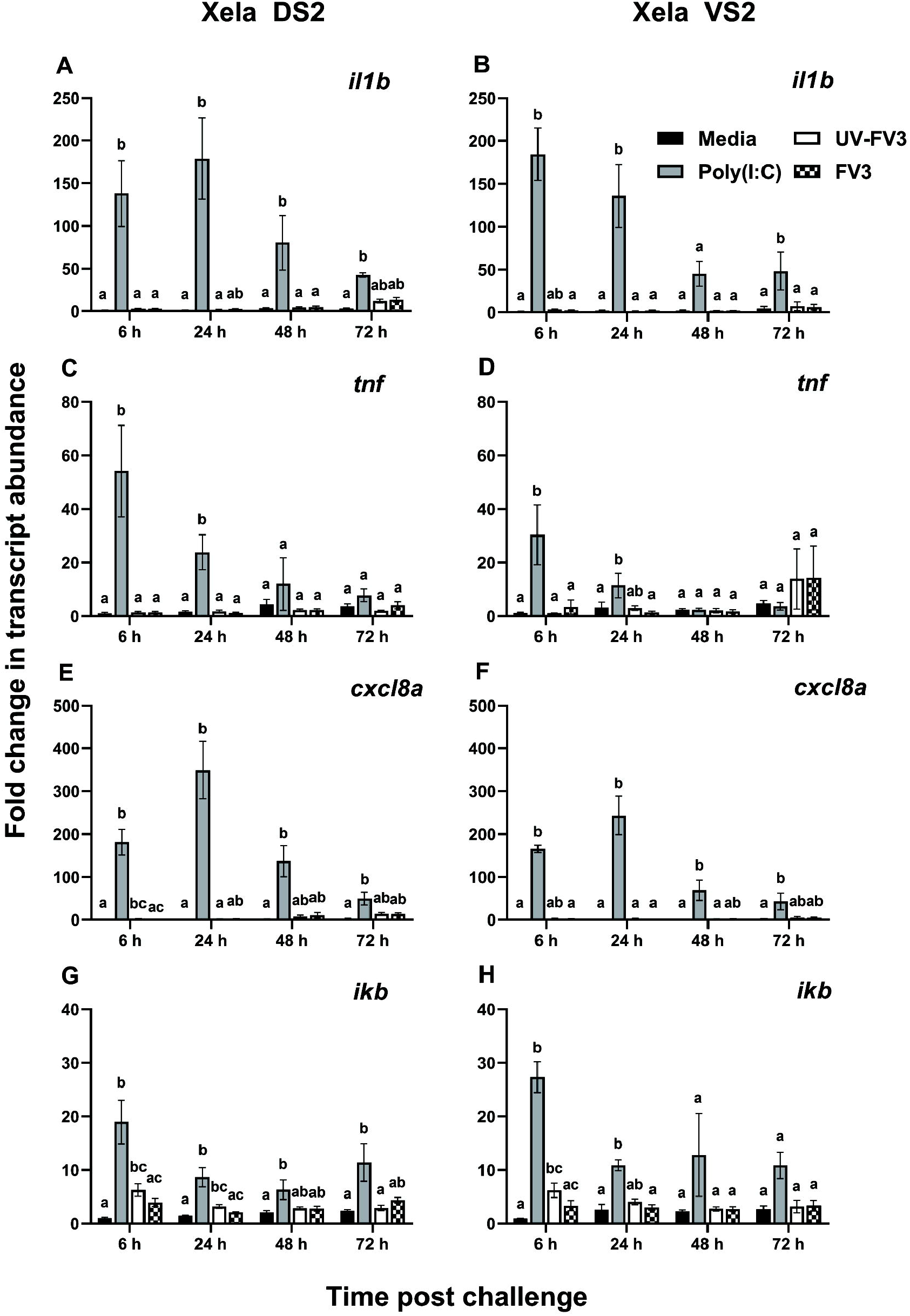
Xela DS2 and Xela VS2 do not upregulate proinflammatory gene transcripts following challenge with UV-inactivated FV3 or FV3. Xela DS2 or Xela VS2 were mock-infected (media), treated with 1 μg/mL poly I:C (pIC), challenged with UV-inactivated FV3 at a MOI of 2 or infected with FV3 at a MOI of 2 for 6 h, 24 h, 48 h or 72h. RT-qPCR was performed using cDNA generated from each treatment and time point to determine relative transcript levels of (A,B) *il1b*, (C,D) *tnf*, (E,F) *cxcl8a*, and (G,H) *ikb*. Data were analyzed using the ΔΔCt method and were expressed as a fold-change in transcript levels relative to that of the 6 h non-treated control sample. Data represent the mean ± standard error and were analyzed with a Kruskal-Wallis test and Dunn’s post-hoc test. *n* = 4 independent experiments. Within each time point, significant statistical differences (*p* < 0.05) are denoted using a lettering system, wherein groups with the same lettering are not statistically different.

### 3.4. Evaluating poly(I:C) cytotoxicity in Xela DS2 and Xela VS2

To evaluate potential cytotoxicity of poly(I:C), Xela DS2 and Xela VS2 were treated with 0 – 10,000 ng/mL of poly(I:C) and cell metabolic activity and membrane integrity were assessed over 5 days post-treatment using a combined resazurin/CFDA-AM assay. The two highest doses of poly(I:C), 1,000 ng/mL and 10,000 ng/mL, induced a significant loss in Xela DS2 cell metabolic activity (Fig. 7A) and membrane integrity (Fig. 7C), and a significant loss in Xela VS2 cell metabolic activity (Fig. 7B) and membrane integrity (Fig. 7D) at virtually all time points examined. No statistically significant differences in cell metabolic activity or cell membrane integrity were observed in Xela DS2 or Xela VS2 across all days when cells were treated with 10, 50 or 100 ng/mL poly(I:C) compared to the non-treated, time-matched controls (Fig. 7). Due to the decrease in Xela DS2 and Xela VS2 cell viability in the presence of high concentrations of poly(I:C), poly(I:C) concentrations of 100 ng/mL or less were used in subsequent experiments.

**Figure 7.**
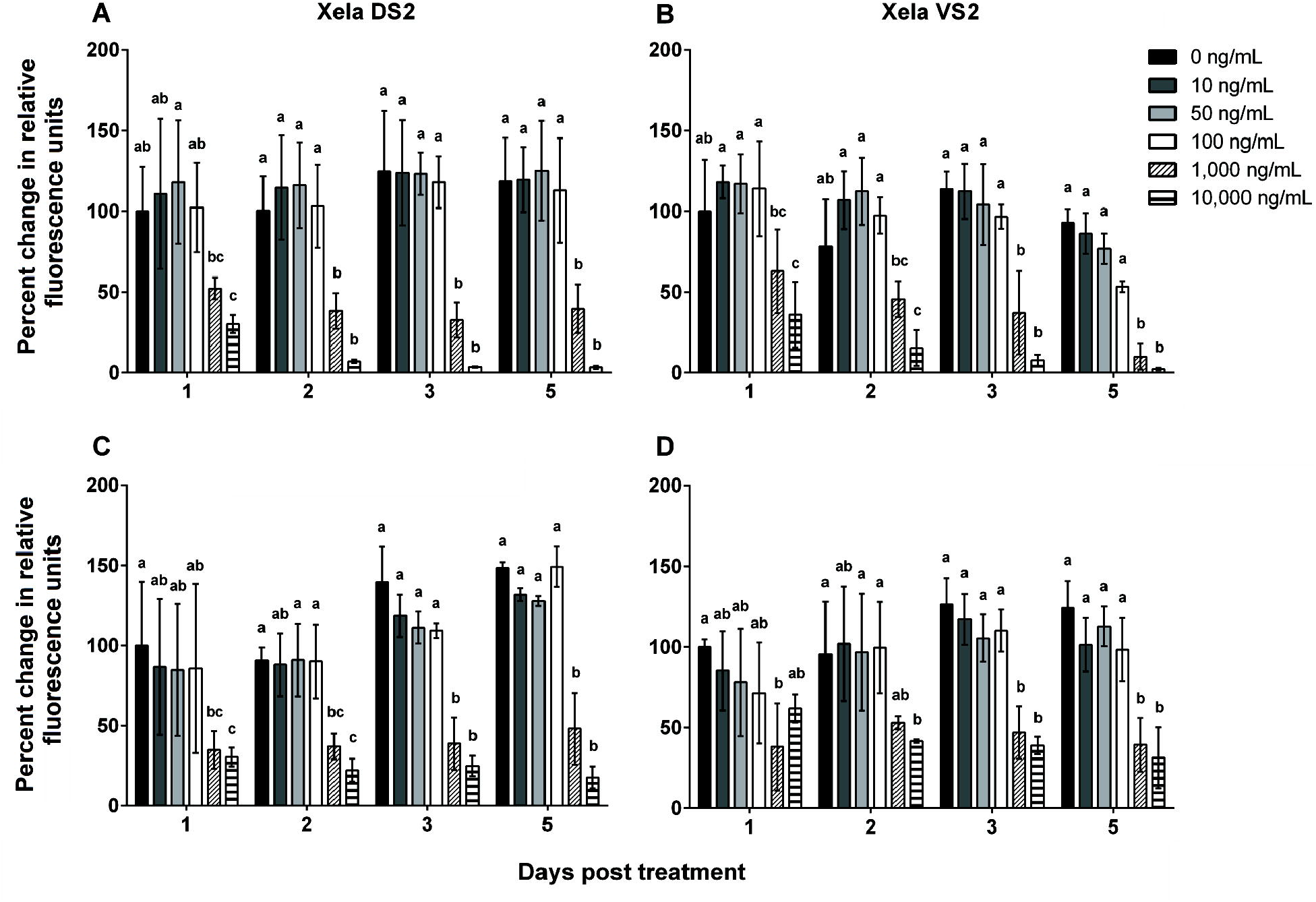
High concentrations of poly(I:C) are cytotoxic to Xela DS2 and Xela VS2. Xela DS2 and Xela VS2 were treated with 0, 10, 50, 100, 1,000 or 10,000 ng/mL poly(I:C) for 1, 2, 3 and 5 days. Cell metabolic activity for (A) Xela DS2 and (B) Xela VS2 was assessed using a resazurin assay and cell membrane integrity for (C) Xela DS2 and (D) Xela VS2 was assessed with CFDA-AM assay. Data are expressed as percent change in fluorescence intensity relative to that of 0 ng/mL poly(I:C) control group at day 1 post-treatment for each cell line and assay. Data represent the mean ± standard error. *n* = 3 independent experiments. Data were analyzed with a two-way ANOVA and Tukey’s post-hoc test. Within each time point, significant statistical differences (*p* < 0.05) are denoted using a lettering system, wherein groups with the same lettering are not statistically different.

### 3.5. Poly(I:C) pretreatment confers partial protection against FV3-induced CPE in Xela DS2 and Xela VS2

To assess the potential effects of poly(I:C) pretreatment on susceptibility of Xela DS2 and Xela VS2 to FV3, cells were pre-treated with 0, 10, 50 or 100 ng/mL of poly(I:C) for 24 h before mock-infection or FV3 infection (MOI 2). To quantify FV3-induced loss of cellular adherence, adherent Xela DS2 (Fig. 8A, Supplementary Figure 5A) and Xela VS2 (Fig. 8B, Supplementary Figure 5B) cell nuclei were enumerated and the numbers of adherent cells in each of the FV3-infected groups were expressed as percentages relative to the number of adherent cells in the corresponding pre-treated, mock-FV3 infected group. Two-way ANOVA analyses indicated a significant effect of time (*p* < 0.0001) and poly(I:C) pretreatment concentration (*p* < 0.0001), accompanied with a significant interaction between these two factors (*p* < 0.0001), on both Xela DS2 (Fig. 8A) and Xela VS2 (Fig. 8B) cell adherence. At 0 dpi and 3 dpi, no statistically significant differences in adherent cell numbers were observed in FV3-infected Xela DS2 (Fig. 8A) or Xela VS2 (Fig. 8B) across any of the poly(I:C) pre-treated groups. By 5 dpi, we observed significant CPE characterized by a loss of cellular adherence in FV3-infected Xela DS2 pre-treated with 0 or 10 ng/mL of poly(I:C) (Fig. 8A, Supplementary Fig. 5A). Meanwhile, pretreatment of Xela DS2 with 50 or 100 ng/mL of poly(I:C) pretreatment appeared to completely mitigate FV3-induced CPE as determined by enumerating adherent cell nuclei (Fig. 8A, Supplementary Figure 5A) and inspection of phase contrast images (Supplementary Figure 6A). In Xela VS2, poly(I:C) pretreatment was able to mitigate FV3-induced CPE in a dose-dependent manner with the 50 and 100 ng/mL poly(I:C) pretreatments conferring partial and complete mitigation of FV3-induced CPE, respectively (Fig. 8B, Supplementary Figure 5B, Supplementary Figure 6B). Cellular adherence of Xela DS2 (Supplementary Figure 5A) or Xela VS2 (Supplementary Figure 5B) was not affected in any of the poly(I:C)-treated mock-infected groups over the course of the experiment.

**Figure 8.**
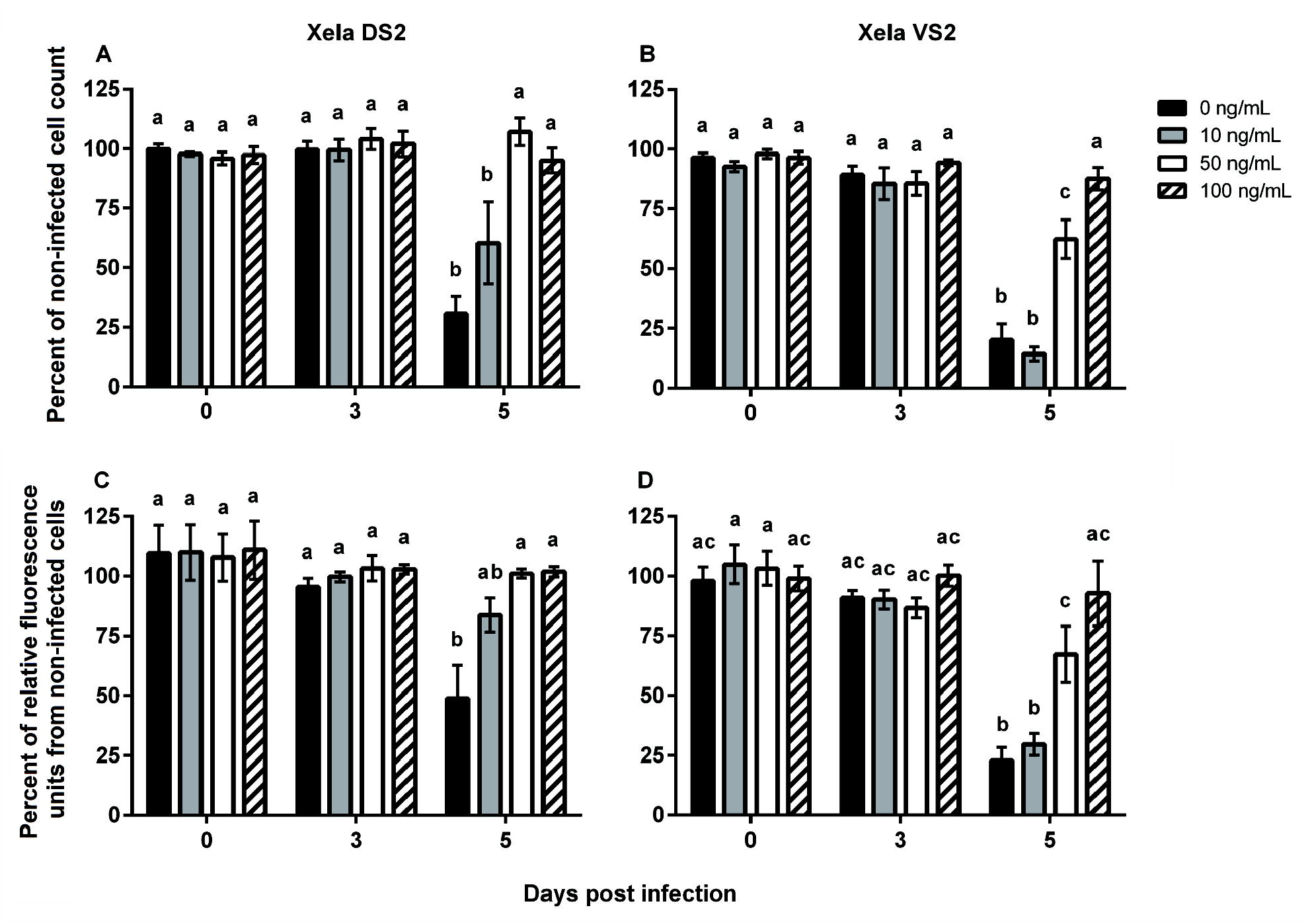
Pretreatment of Xela DS2 and Xela VS2 with poly(I:C) mitigates FV3-induced loss of cell adherence. Xela DS2 and Xela VS2 were pre-treated with 0, 10, 50, or 100 ng/mL poly(I:C) for 24 h before mock-infection or FV3 infection (MOI 2) of cells by absorption for 2 h. On 0, 3, and 5 dpi, cell culture media was removed and the number of adherent (A) Xela DS2 and (B) Xela VS2 cells was determined by enumerating NucBlue Live-stained nuclei using the fluorescent imaging and cell count feature of the BioTek Cytation 5 Gen5 software. Cell viability of adherent (C) Xela DS2 and (D) Xela VS2 was assessed using the metabolic-based resazurin assay. Data represent the mean ± standard error. *n* = 3 independent experiments. Data were analyzed with a two-way ANOVA and Tukey’s post-hoc test. Significant statistical differences (*p* < 0.05) across all groups are denoted using a lettering system, wherein groups with the same lettering are not statistically different.

Similar mitigation of FV3-induced CPE was observed in poly(I:C) pre-treated FV3-infected Xela DS2 (Fig. 8C) and Xela VS2 (Fig. 8D) as assessed through a cell metabolic assay as a proxy for cell viability. Two-way ANOVA analysis indicated that there was a significant effect due to time (*p* = 0.0004 for Xela DS2, *p* < 0.0001 for Xela VS2) and poly(I:C) pretreatment concentration (*p* = 0.0159 for Xela DS2, *p* = 0.0002 for Xela VS2), accompanied with a significant interaction between these two factors (*p* = 0.0308 for Xela DS2, *p* < 0.0001 for Xela VS2), on Xela DS2 (Fig. 8C) and Xela VS2 (Fig. 8D) cell metabolic activity. Pretreatment of Xela DS2 with 50 or 100 ng/mL of poly(I:C), but not 10 ng/mL, was able to abrogate FV3-induced CPE (Fig. 8C), while pretreatment of Xela VS2 with 50 or 100 ng/mL poly(I:C) provided partial or complete abrogation of FV3-induced CPE, respectively (Fig. 8D).

### 3.6. Poly(I:C) pretreatment limits FV3 replication in Xela DS2 and Xela VS2

To determine whether poly(I:C) pretreatment limited FV3 replication, Xela DS2 or Xela VS2 were pre-treated with low serum media alone [0 ng/mL poly(I:C)] or containing 100 ng/mL poly(I:C) for 24 h prior to their infection with FV3. TCID_50_/mL values were determined for virus-containing cell culture media collected from 0, 3 and 5 dpi FV3-infected Xela DS2 or Xela VS2 cells that had been pre-treated with either 0 ng/mL or 100 ng/mL of poly(I:C) (Fig. 9). Two-way ANOVA analysis indicated a significant effect of time (Xela DS2, Xela VS2, *p* < 0.0001) and poly(I:C) pretreatment (Xela DS2, *p* < 0.0001; Xela VS2, *p* = 0.0106), accompanied with a significant interaction between these two factors (Xela DS2, *p* < 0.0001; Xela VS2, *p* = 0.0063), on Xela DS2 and Xela VS2 TCID_50_/mL values. Poly(I:C) pretreatment at 100 ng/mL limited FV3 replication in both Xela DS2 (Fig. 9A) and Xela VS2 (Fig. 9B) compared to non-pretreated cells that showed a time-dependent increase in TCID_50_/mL values. Although cell culture media from poly(I:C) pretreated Xela DS2 had higher TCID_50_/mL values at 3 dpi compared to 0 dpi, there was a significant reduction in TCID_50_/mL values compared to the non-treated infected control at 3 dpi. By 5 dpi, the TCID_50_/mL values of cell culture media from poly(I:C) pretreated Xela DS2 were significantly lower than the time-matched infected group, to the extent that these values were not statistically different from either of the 0 dpi treatment groups (Fig. 9A). In contrast, poly(I:C)-pretreated Xela VS2 did not appear to limit FV3 TCID_50_/mL values to the same extent, with a significant reduction in TCID_50_/mL values only observed at 5 dpi in the cell culture media of the poly(I:C) pre-treated group relative to the non-treated group (Fig. 9B).

**Figure 9.**
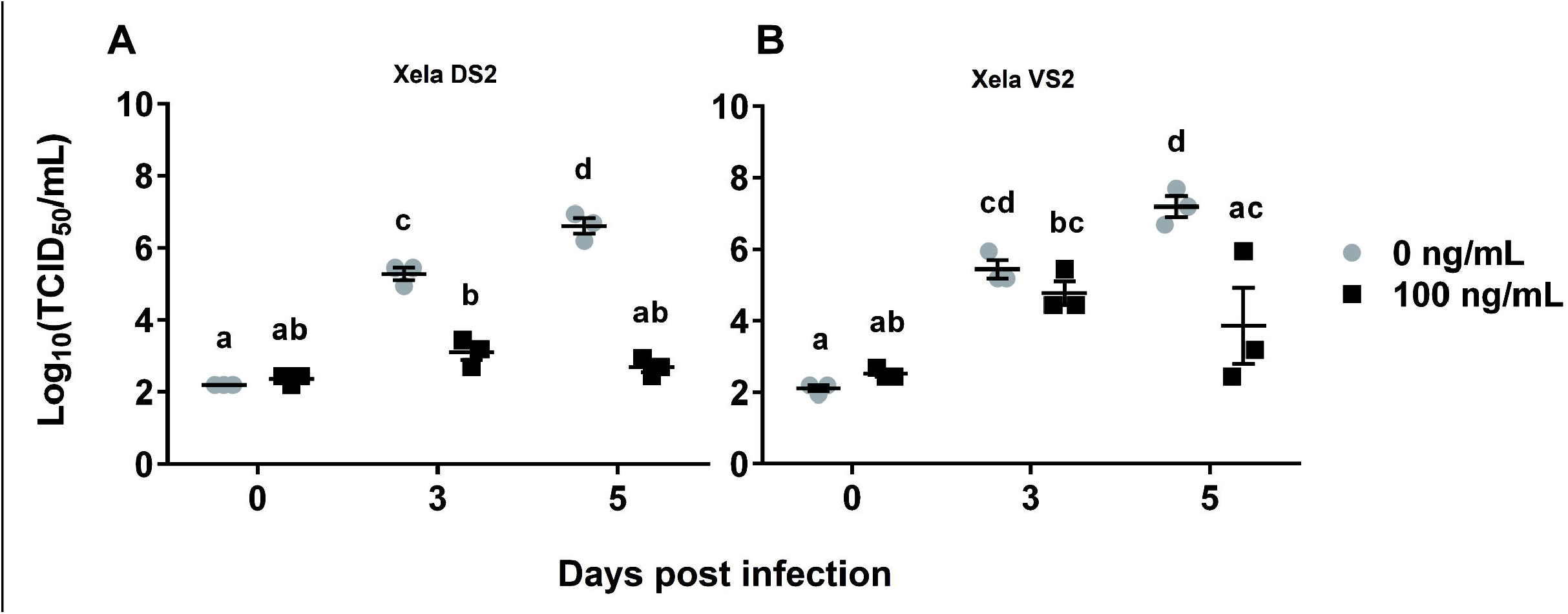
Poly(I:C) pretreatment limits FV3 replication in Xela DS2 and Xela VS2. Xela DS2 and Xela DS2 were pre-treated with 0 or 100 ng/mL poly(I:C) for 24 h before FV3 infection (MOI 2) of cells by absorption for 2 h. Following pretreatment and infection, virus-containing media was collected from (A) Xela DS2 and (B) Xela VS2 at 0, 3, and 5 dpi and used to determine TCID_50_/mL values. Data were log10 transformed and are presented as the mean ± standard error of three independent experiments. Data were analyzed with a two-way ANOVA and Tukey’s post-hoc test. Significant statistical differences (*p* < 0.05) are denoted using a lettering system, wherein groups with the same lettering are not statistically different.

## 4. Discussion

As amphibian skin epithelial cells are the first cellular line of defence against invading pathogens, examining the initial response of these cells to FV3 is essential to furthering our understanding of FV3 pathogenesis. Our previous research resulted in the successful expansion of the *X. laevis* invitrome to include two novel skin-epithelial cell lines (Xela DS2 and Xela VS2) that demonstrate the ability to initiate antiviral and proinflammatory gene regulatory responses upon treatment with poly(I:C), a known inducer of type I IFNs (Bui-Marinos et al. 2020). In this study, we utilized Xela DS2 and Xela VS2 as a novel platform to evaluate frog skin epithelial cell permissibility to FV3, characterize frog skin epithelial cell antiviral and proinflammatory gene regulatory responses to FV3 and UV-inactivated FV3, and demonstrate prior establishment of antiviral programs in these cell lines confers protection against FV3 infection.

Xela DS2 and Xela VS2 skin epithelial-like cells appear equally permissive to FV3 and exhibit cell rounding and detachment CPE similar to that observed in other FV3-permissive cell lines (Chinchar et al., 2003; Pham et al., 2015). Xela DS2 and Xela VS2 produce relatively high levels of infectious FV3 virions [log_10_(TCID_50_/mL) of 6-7], albeit not to the same levels observed in the EPC cell line [log_10_(TCID_50_/mL) of 8-9] (Pham et al., 2015). However, Xela DS2 and Xela VS2 monolayers do not achieve the same cell density as EPC monolayers and the apparent lower production of infective FV3 particles may be a result of fewer host cells to act as viral factories. Interestingly, while Xela DS2 and Xela VS2 are permissive to FV3 at higher MOIs (> 0.02), CPE and increases in TCID_50_/mL values were not observed when these cell lines were infected with FV3 at a MOI of 0.002 over the seven days examined, even though low levels of viral particles were detected. It is possible this lack of viral replication at a low MOI is due to ineffective binding of viral particles or some level of cell-intrinsic viral restriction in Xela DS2 and Xela VS2. Our findings support the use of Xela DS2 and Xela VS2 to propagate FV3 and for use as *in vitro* models in which interactions between frog skin epithelial cells and FV3 can be examined.

FV3 exists as enveloped and non-enveloped/naked virions and is known to enter host cells through receptor-mediated endocytosis (enveloped virions) or fusion at the plasma membrane followed by nucleocapsid injection into the cytoplasm (naked virions) (Braunwald et al., 1985; Gendrault et al., 1981; Houts et al., 1974; Kelly, 1975). In contrast to previous studies that demonstrated that FV3 uses class A scavenger receptors for cellular entry in tadpole cell lines (American toad, wood frog, green frog, bullfrog) and adult *X. laevis* macrophages (Vo et al., 2019a), Xela DS2 and Xela VS2 do not express appreciable levels of class A scavenger receptor (*srai/ii*, *scara3*, *scara4*, *scara5*, and *marco*) transcripts despite being susceptible and permissive to FV3. Although we did not examine Xela DS2 and Xela VS2 for the presence of class A scavenger receptor proteins, our data suggests that either FV3 is utilizing different cell-surface receptors for endocytosis-mediated entry, or that primarily naked FV3 virions are entering Xela DS2 and Xela VS2. Given the broad cell and tissue tropism of FV3, it is likely that additional host cell receptors are utilized by FV3 to gain entry to diverse cell types/tissues and warrants further study.

As Xela DS2 and Xela VS2 are derived from *X. laevis* skin and are epithelial-like, these cell lines serve as ideal *in vitro* systems to expand the arsenal of *X. laevis* resources at our disposal to investigate frog skin epithelial antiviral defences against FV3. Although susceptible to FV3, Xela DS2 and Xela VS2 failed to induce the expression of the key antiviral and proinflammatory transcripts measured herein when infected with FV3 or UV-inactivated FV3. Our observations agree with the current theory that FV3 encodes a capsid protein that is responsible for the immediate immunoevasion of host antiviral pathways independent of viral transcription (Robert et al., 2017; Rothenburg et al., 2011). However, our findings contrast with observations of induction of innate antiviral pathways, such as type I IFN transcripts, in response to FV3 infection of adult *X. laevis* skin tissue *in vivo* and adult *X. laevis* skin cells (mixed cell type populations isolated from whole tissues) *in vitro* (Wendel et al., 2017; Wendel et al., 2018). We propose that the interactions between skin epithelial cells and macrophages (or perhaps other cell types) present in skin tissues may explain this discrepancy. In *X. laevis*, macrophages have been identified as important mediators of FV3 infection (Grayfer and Robert, 2014) and distinct differentiated macrophage subpopulations have been demonstrated to condition cell media with type I IFNs that can be used to confer resistance to FV3 in the susceptible *X. laevis* kidney epithelial A6 cell line (Yaparla et al., 2018). As macrophages have been found to reside in frog skin (Fox and Whitear, 1990; Lehman, 1953), we hypothesize that initial FV3 replication in skin epithelial cells and subsequent cell death leads to the release of viral material (e.g. apoptotic bodies; viral RNA through cell necrosis) that is phagocytosed by resident and/or recruited macrophages. Activation of antiviral programs in macrophages following the detection of exogenous viral pathogen-associated molecular patterns (PAMPs) by endosomal pattern recognition receptors may account for the detection of type I IFN transcripts observed in frog skin tissues or in mixed cell type populations. Secretion of type I IFN by activated macrophages would initiate antiviral programs in neighbouring skin cells, thus potentially conferring protection to FV3 replication in skin tissues. Collectively, these observations highlight the importance of studying the contributions of individual cell types and as well as cellular interactions within complex skin tissue environments.

Simulation of Xela DS2 or Xela VS2 with higher concentrations of extracellular poly(I:C) [10 μg/mL, (Bui-Marinos et al., 2020); 1 μg/mL, this study] results in a modest induction of antiviral genes and a robust upregulation of proinflammatory cytokine genes, strongly supporting that idea that frog skin epithelial cells are important cellular participants in mediating skin antiviral defences, in addition to their role as a physical barrier. Aside from regulating antiviral and proinflammatory genes, concentrations of poly(I:C) above a threshold (> 100 ng/mL) are cytotoxic to Xela DS2 and Xela VS2. Similar poly(I:C)-induced cytotoxicity has been noted in an *Anaxyrus americanus* tadpole cell line (Vo et al., 2019b), yet many other vertebrate cells do not exhibit poly(I:C)-induced cytotoxicity at these concentrations [e.g. (Kumar et al., 2006; Ritter et al., 2005)]. Our findings suggest that an increased sensitivity of frog skin epithelial cells to viral nucleic acids may be advantageous for removal of infected cells to limit viral spread and, together with the induction of effector proinflammatory cytokines and chemokines, is an important antiviral defence mechanism of frog skin epithelial cells.

To ascertain frog skin epithelial cell immunocompetence in relation to functional viral restriction to FV3, we prestimulated Xela DS2 and Xela VS2 with poly(I:C) concentrations below the observed threshold for poly(I:C)-induced cytotoxicity (100 ng/mL or less) prior to FV3 infection. Consistent with previous observations of reductions in FV3 replication following pretreatment of *X. laevis* A6 cells with type I IFN (Grayfer et al., 2014), subcutaneous injection of *X. laevis* tadpoles with type I IFN (Wendel et al., 2017), or pretreatment of rainbow trout gonadal fibroblast cell lines with poly(I:C) (Lisser et al., 2017), pretreatment of Xela DS2 and Xela VS2 with low concentrations of poly(I:C) mitigated FV3-induced CPE and limited FV3 replication. It is likely that these antiviral effects are the result of the induction of type I IFN-mediated antiviral programs, as poly(I:C) is a well-known inducer of type I IFN and we have shown that poly(I:C) induces the expression of antiviral and proinflammatory gene transcripts in these cell lines [this study and (Bui-Marinos et al., 2020)]. Interestingly, the induction of functional anti-FV3 restrictions and its protective effects appear stronger in Xela DS2 than Xela VS2, as immune gene transcripts are generally induced earlier in response to poly(I:C) treatment [this study and (Bui-Marinos et al., 2020)] and protection against FV3 replication is achieved at lower poly(I:C) doses, and to a greater extent, in Xela DS2 compared to Xela VS2. Whether the differential induction of poly(I:C) mediated anti-FV3 responses observed between Xela DS2 and Xela VS2 are reflective of physiological differences between skin epithelial cells from the dorsal and ventral skin, as well as the underlying mechanism(s) driving these differences, are not known and require further study.

Our results provide evidence of frog skin epithelial cell immunocompetence in establishing a functional antiviral state. Additionally, we observed that prior induction of this antiviral state with poly(I:C) is effective against FV3 immunoevasion mechanisms, thereby highlighting these cells as important contributors to skin innate immune defences. In frog skin, induction of an antiviral state in epithelial cells may require paracrine type I IFN signaling from resident immune cells (e.g. macrophages) or other cell types within the complex skin tissue to initiate an effective anti-FV3 state. Indeed, the restricted tissue necrosis observed in infected frogs supports the eventual establishment of an antiviral state in frog skin tissues. The ability of frog skin epithelial cells to detect and respond to a synthetic viral PAMP analogue suggests that Xela DS2 and Xela VS2 cell lines can be used to identify and/or screen immunomodulatory molecules present in the host or the environment that may impact epithelial cell susceptibility/permissibility to FV3. Thus, in addition to furthering our understanding of antiviral responses of skin epithelial cells to FV3 infection, we believe Xela DS2 and Xela VS2 represent novel *in vitro* platforms for future research in frog epithelial cell-FV3 interactions and immunoevasion mechanisms at the host-environment interface.

## 6. Author contributions

**Maxwell P. Bui-Marinos:** conceptualization, methodology, investigation, data curation, formal analysis, writing – original draft, review and editing. **Lauren A. Todd**: data curation, formal analysis, investigation, methodology, visualization, writing - original draft, review and editing. **Marie-Claire D. Wasson**: investigation, writing - review and editing. **Brandon E.E. Morningstar**: investigation, formal analysis, writing - review and editing. **Barbara A. Katzenback**: conceptualization, methodology, investigation, resources, writing – original draft and review and editing, supervision, funding acquisition.

## 7. Acknowledgements

This work was supported by a Natural Sciences and Engineering Research Council (NSERC) Discovery Grant (RGPIN-2017-04218) and University of Waterloo Start-Up Funds to BAK, University of Waterloo Science Graduate Awards to MPB-M, and a NSERC Postdoctoral Fellowship (PDF-546075-2020) to LAT. We thank Dr. Niels C. Bols for providing the FV3 strain and EPC cell line used in this study and for kindly permitting us to use their cell culture facilities during the early part of this study, Dr. Nguyen T. K. Vo for providing technical training on FV3 infection procedures and maintenance of the EPC cell line, and Adrienne Ranger for their technical assistance in initial testing of scavenger receptor primers.

