## Supplemental Methods for "Prior induction of cellular antiviral pathways limits frog virus 3 replication in two permissive *Xenopus laevis* skin epithelial-like cell lines"

### Supplementary Methods

#### 1. Morphological changes in early passage Xela DS2 and Xela VS2 after FV3 challenge

Xela DS2 (passage 23) and Xela VS2 (passage 25) ($n$ = 1) were seeded in 25 $cm^2$ plug-seal tissue-culture treated flasks (Thermo Fisher Scientific) at a final cell density of $6 \times 10^6$ cells/flask in 2 mL of Xela complete media and allowed to adhere overnight at 26 °C. The next day, Xela complete media was removed from flasks and cells were treated with 2 mL of Xela low serum media alone (mock-infected control), or containing FV3 at MOIs of 0.2, 2 or 20. After 2 h of incubation at 26 °C, media was removed and wells were washed 3 times with 2 mL APBS, prior to the addition of 2 mL fresh Xela low serum media to all flasks. At 24 hpi, cell morphology was examined by capturing phase contrast digital images using a Nikon Eclipse TSX-100 fitted with a color camera and Picture Project software.

#### 2. Susceptibility of early passage Xela DS2 and Xela VS2 to FV3

Xela DS2 and Xela VS2 cells (passages 20-55 were seeded in 48-well tissue-culture treated plates (BioLite, Thermo Scientific) at a final cell density of 50,000 cells/well in 0.2 mL Xela complete media and allowed to adhere overnight at 26 °C. The next day, media was removed and cells were treated with 0.25 mL Xela low serum media alone (mock-infected control), or containing FV3 at a MOI of 0.002, 0.02, 0.2, 2 or 20 in triplicate wells. After 2 h of incubation at 26 °C, media was removed and wells were washed three times with 300 μL APBS before the addition of 500 μL Xela low serum media. FV3-induced CPE was evaluated using Alamar Blue (Life Technologies Inc) and is a commercial preparation of resazurin (Rampersad, 2012). Cell metabolic activity was assessed using Alamar Blue at 1, 3, 5 7, 10 and 14 dpi. Triplicate wells containing the same volume of virus-free media without cells in the well were used to determine background levels of fluorescence and the average fluorescence of these wells were subtracted from all other values. Cell culture media was removed prior to the addition of 200 μL Alamar Blue solution (diluted 1:20 with APBS) to each well. Plates were incubated for 1 h in the dark and read on a BioTek Synergy H1 plate reader (excitation wavelength = 535 nm and emission wavelength = 590 nm). Data are expressed as relative fluorescent units. This experiment was performed seven independent times using cells from different passages ($n$ = 7).

#### 3. Permissibility of early passage Xela DS2 and Xela VS2 to FV3

Xela DS2 (passages 30 – 50) or Xela VS2 (passages 30 – 50,) were seeded in 12-well plates (BioLite, Thermo Scientific) at a final cell density of 250,000 cells/well in 0.75 mL complete media and allowed to adhere overnight at 26 °C. The next day, complete media was removed from wells and cells treated with 0.5 mL of low serum media containing FV3 at a MOI of 0.2 or 20. After 2 h of incubation at 26°C, media was removed and wells were washed three times with 0.5 mL APBS, followed by the addition of 2.5 mL low serum media to all wells. Cell culture media was collected at 0 (30 min post-wash), 1, 3, and 5 dpi for FV3 challenge at a MOI of 20, and at 0, 1, 3, 5, 7, 10, and 14 dpi for FV3 challenge at a MOI of 0.2. Collected cell culture media was centrifuged at 500 × $g$ for 5 min and supernatants were transferred to a sterile 1.5 mL microcentrifuge tube prior to storage at -80 °C. This experiment was performed three independent times using cells from different passages ($n$ = 3). Determination of $TCID_{50}$/mL values were performed as described in Section 2.4.

#### 4. Detection of FV3 *mcp* transcripts in FV3-infected Xela DS2 and Xela VS2

Xela DS2 (passages 75 – 90) and Xela VS2 (passages 75 – 90) cells were seeded in 6-well tissue-culture treated plates at a final cell density of 625,000 cells/well in 1 mL Xela complete media and allowed to adhere overnight at 26 °C. The next day, cells were treated with 6.25 mL low serum media alone (mock-infected control), or containing FV3 at MOIs of 0.2 or 20, for 2 h at 26 °C. Afterwards, media was removed and wells were washed three times with 1 mL APBS prior to the addition of 2 mL fresh Xela low serum media. Plates were then incubated at 26 °C for the duration of the experiment. Cells were collected at 12 and 24 hpi for cells challenged with FV3 at a MOI of 20, or 1 and 3 dpi for cells challenged with FV3 at a MOI of 0.2. Cell pellets were resuspended in with 500 µL APBS and centrifuged at 500 × $g$ for 5 min to remove trace serum. RNA was isolated from the cell pellets using EZ-10 Spin Column Total RNA Minipreps Super Kit (Section 2.7) and cDNA was synthesized using the SensiFAST cDNA Synthesis Kit (BioLine) according to the manufacturer's specifications. Briefly, 500 ng of RNA was mixed with 1 U of reverse transcriptase and 4 µL of 5 × reaction buffer in a total volume of 20 µL. The reactions were then incubated at 25 °C for 10 min, 42 °C for 15 min and inactivated at 85 °C for 5 min. RT-PCR reactions consisted of 2 mM $MgCl_2$ (GeneDireX), 200 μM dNTP, 200 nM sense primer (Sigma), 200 nM antisense primer (Sigma), 0.625 U Taq DNA Polymerase (GeneDireX), and 0.5 μL template in 25 μL reaction volumes. Targets included *mcp* viral transcripts from FV3 (Sense: 5'-GACTTGGCCACTTATGAC-3', Antisense: 5'-GTCTCTGGAGAAGAAGAA-3') (Pham et al., 2015), and *X. laevis actb* (Sense: 5'-AGGAGATGAAGCTCAAAGCAA-3', Antisense: 5'-GTTACACCATCACCTGAGTCC-3') as an endogenous control. Thermocycling conditions were as follows: initial denaturation at 95 °C, 5 min; 25 (*actb*) or 35 (*mcp*) amplification cycles of denaturation at 95 °C for 45 s, annealing at 55 °C for 30 s, and elongation at 72 °C for 45 s (*actb*) or 60 s (*mcp*), followed by a final extension step at 72 °C for 10 min. To each RT-PCR reaction, 5 μL of 6× loading buffer [0.1% xylene cyanol (ICN Biomedicals), 30% v/v glycerol (EMD Chemicals)] was added prior to loading into a 1.4% agarose (VWR) gel containing 1× RedSafe Nucleic Acid Staining Solution (FroggaBio) and electrophoresed in 1× TAE buffer at 140 V for 20 min. Gels were imaged using a ChemiDoc imager (BioRad) with the Image Lab program. A representative experiment of three independent experiments is shown.

### 5. Confirmation of UV-inactivation of FV3

To verify UV inactivation of FV3, viral *mcp* transcripts and $TCID_{50}$/mL values were measured in EPC cells treated with FV3 or UV-FV3 ($n = 1$). EPC cells were seeded in a 6-well plate (Thermo Fisher Scientific) at a final cell density of 800,000 cells/well in 1 mL EPC complete media and allowed to adhere overnight at 26 °C. After 48 h, media was removed and cells treated with 1 mL EPC low serum media alone (mock-infected control), UV-inactivated FV3 (MOI of 2) or FV3 (MOI of 2) in triplicate, for 2 h at 26 °C. Afterwards, treatment-containing media was removed from EPC monolayers and wells were washed three times with 1 mL of APBS prior to the addition of 2 mL fresh EPC low serum media. Plates were incubated at 26 °C for the duration of the experiment. At each time point, media was collected for use in determining $TCID_{50}$/mL values (Section 2.4) and cells were collected for use in total RNA isolation (Section 2.7) and cDNA synthesis (Supplementary Methods Section 4). FV3 *mcp* transcripts were detected as described in Supplementary Methods Section 4, and EPC *actb* (Sense: 5'-

TGAAGATCCTGACCGAGAGA-3', Antisense: 5'-GGATACCGCAAGACTCCATAC-3') transcripts were amplified using the following thermocycling conditions: initial denaturation at 95 °C, 5 min; 30 amplification cycles of denaturation at 95 °C for 45 s, annealing at 55 °C for 30 s, and elongation at 72 °C for 50 s, followed by a final extension step at 72 °C for 10 min. Amplified products were electrophoresed and visualized on an agarose gel as described in Supplementary Methods Section 4.

### 6. Measurement of FV3 *mcp* transcripts and viral titres in Xela DS2 and Xela VS2 treated with poly(I:C), UV-inactivated FV3, and FV3

The treatment of Xela DS2 and Xela VS2 was performed as described in Section 2.9. Cells were collected for total RNA isolation and cDNA synthesis (Section 2.9 and 2.10) and the resulting cDNA used as a template in RT-PCR to detect FV3 *mcp* transcripts in Xela DS2, Xela VS2, and EPC cells as described in Supplementary Methods Sections 4 and 5. Cell culture media was collected from all treatments and used to determine $TCID_{50}$/mL values (Section 2.4).
