## Supplemental Figure Legends for "Prior induction of cellular antiviral pathways limits frog virus 3 replication in two permissive *Xenopus laevis* skin epithelial-like cell lines"

**Supplementary Figure 1. Effect of FV3 challenge on early passage Xela DS2 and Xela VS2**. (A) Phase contrast images of early passage Xela DS2 and Xela VS2 24 h after challenge with a mock-infected control or with FV3 at MOI of 0.2, 2, or 20. Images were taken at 50× magnification using a Nikon Eclipse TSX-100 microscope (scale bar = 500 μm). Images shown are from one representative trial. (B) Early passages of Xela DS2 and Xela VS2 were infected with FV3 at a MOI of 0.0002, 0.002, 0.02, 0.2, 2, or 20 and metabolic activity of adherent cells was monitored over 14 days using an Alamar Blue assay and a BioTek Synergy H1 plate reader. Data is expressed as relative fluorescent units and represent the mean ± standard error ($n$ = 7 independent experiments). Significant differences were determined by a Kruskal-Wallis test and a Dunn's post-hoc test ($p < 0.05$), wherein asterisks indicate significant differences in relation to the corresponding time matched mock-infected (MOI 0) controls. (C) Early passages of Xela DS2 and Xela VS2 were infected with FV3 at a MOI of 0.2 or 20 and cell culture media was collected over 5 or 14 d, respectively. Cell culture media from FV3-infected Xela DS2 and Xela VS2 was serially diluted and applied to a 96-well plate containing a monolayer of EPC cells. EPC monolayers were scored for cytopathic effects (CPE) after 7 d to determine $TCID_{50}$/mL values. Dashed lines represent projected $TCID_{50}$/mL values as no CPE was observed at day 0 for Xela DS2 or Xela VS2 cells infected at a MOI of 0.2. CPE were not observed in cell culture media from non-infected Xela DS2 and Xela VS2 cell lines at any time point. Data represent the mean ± standard error, $n$ = 3 independent experiments.

**Supplementary Figure 2. FV3 major capsid protein transcripts are detected in Xela DS2 and Xela VS2 after FV3 infection.** RT-PCR analysis of FV3 major capsid protein (*mcp*) transcripts in no template control (NTC), Xela DS2 and Xela VS2 mock-infected controls (C) and FV3 infected (I) cells at 12 and 24 hpi (MOI 20) or at 24 and 72 hpi (MOI 0.2). Detection of *actb* transcripts was used as an endogenous control.

**Supplementary Figure 3. Xela DS2 and Xela VS2 lose adherence when treated with poly(I:C), UV-inactivated FV3, or FV3.** Phase-contrast images of Xela DS2 (A) and Xela VS2 (B) cells when treated with media alone (mock-infected control), 1 μg/mL poly(I:C), or UV-inactivated FV3 or FV3 at a MOI of 2. Cellular morphology was monitored at 0, 6, 24, 48, and 72 hpi by capturing phase contrast images. Images were taken at 200× magnification using a Leica DMi1 microscope (scale bar = 100 μm). Images shown are representative of four independent trials.

**Supplementary Figure 4. Confirmation of UV-FV3 inactivation via RT-PCR and viral titre analysis.** (A) RT-PCR analysis of FV3 *mcp* transcripts in no template control (NTC), Xela DS2 and Xela VS2 mock-infected control (M), poly(I:C)-treated (P), UV-FV3 challenged (MOI 2; ΔF) and FV3-infected (MOI 2; F) cells at 48 and 72 hpi. EPC cells infected with FV3 at a MOI of 2 was used as a positive control. Cell culture media was collected across all treatments at all time points for Xela DS2 (B) and Xela VS2 (C), serially diluted, applied to EPC monolayers and scored for cytopathic effects (CPE) after 7 days to determine the $TCID_{50}$/mL values. CPE were not observed in cell culture media from non-infected or poly(I:C)-stimulated Xela DS2 and Xela VS2 cell lines at any time point, with minimal cytopathic effects observed in UV-FV3-challenged cells. Data represent the mean ± standard error, $n = 4$ independent experiments. Significant differences were determined by a one-way ANOVA and a Tukey's post-hoc test ($p < 0.05$), wherein differential lettering indicates statistical significance between timepoints within a treatment group.

**Supplementary Figure 5. Pre-treatment of Xela DS2 and Xela VS2 with poly(I:C) mitigates FV3-induced cytopathic effects.** Fluorescence microscopy of NucBlue Live-stained Xela DS2 (A) and Xela VS2 (B) cells pre-treated with 0, 10, 50, or 100 ng/mL poly(I:C) for 24 h followed by FV3 infection at a MOI of 2. Cellular morphology was monitored 0, 3, and 5 dpi. The BioTek Cytation 5 imaging reader was used to capture sixteen 40 × images in a single well, stitched together with Gen5 software (scale bar = 200 μm). Images shown are representative of four independent trials.

**Supplementary Figure 6. Monitoring Xela DS2 and Xela VS2 morphology pre-treated with poly(I:C) and subsequently infected with FV3.** Phase-contrast microscopy of Xela DS2 (A) and Xela VS2 (B) cells pre-treated with 0, 10, 50, or 100 ng/mL poly(I:C) for 24 h followed by FV3 infection at a MOI of 2. Cellular morphology was monitored 0, 3, and 5 dpi using the BioTek Cytation 5 imaging reader at 200 × magnification (scale bar = 50 μm). Images shown are representative of four independent trials.
