## Supplementary Table 1 for "Prior induction of cellular antiviral pathways limits frog virus 3 replication in two permissive *Xenopus laevis* skin epithelial-like cell lines"

**Supplementary Table 1.** Primers used for detection of scavenger receptor transcripts

| Target | Sense primer (5' – 3')<br>Antisense primer (5' – 3') | Citation |
|---|---|---|
| *srai/ii* | GAGTCCAGTTCTGTGCAGTTAG<br>GGGCCAGGAAATCCTCTTAATC | This study |
| *scara3* | GTCCAGGAACGGTACGATATTAG<br>ATCGAACGTCATCCAGGTATTT | This study |
| *scara4* | GGATTCAAAGCACGGACAAC<br>CCAGAGGACAACCAGTCACC | This study |
| *scara5* | CACGAGATGGCTCTACTGAATAA<br>GAGAAGCTGCACTGTAGGATAC | This study |
| *marco* | TGGGGTGGTCATCTGCCGAATGTTG<br>GTTGGGTTTCTGACACTGCAGGATACTG | (Vo et al., 2019) |
| *actb* | AGGAGATGAAGCTCAAAGCAA<br>GTTACACCATCACCTGAGTCC | This study |

Vo, N.T.K., Everson, J., Moore, L., DeWitte-Orr, S.J., 2019. Class A scavenger receptor expression and function in eight novel tadpole cell lines from the green frog (*Lithobates clamitans*) and the wood frog (*Lithobates sylvatica*). Cytotechnology, 757-768.
