## Supplementary Table 2 for "Prior induction of cellular antiviral pathways limits frog virus 3 replication in two permissive *Xenopus laevis* skin epithelial-like cell lines"

**Supplementary Table 2.** M-scores of candidate endogenous gene targets used in RT-qPCR

| **Target** | **Xela DS2** | **Xela VS2** |
|---|---|---|
| *actb* | 0.428 | 0.424 |
| *cyp* | 0.393 | 0.440 |
| *ef1a* | 0.479 | 0.600 |
| *gapdh* | 0.434 | 0.462 |
| *hgprt* | 0.462 | 0.534 |
